# Prevalence of Oxalotrophy in the Human Microbiome

**DOI:** 10.1101/2025.03.24.644979

**Authors:** Thomas Junier, Fabio Palmieri, Niki D. Ubags, Aurélien Trompette, Angela Koutsokera, Pilar Junier, Marco Pagni, Samuel Neuenschwander

## Abstract

Incomplete degradation of oxalate, a compound commonly found in the diet, can cause disease in humans, particularly of the kidney. Its concentration in the body depends on several factors, one of which is intestinal absorption, which is itself affected by oxalotrophy among enteric bacteria. Oxalotrophy in the human microbiome is poorly known. In this study, we perform a systematic search for the simultaneous presence of the three oxalotrophy genes, namely *frc*, *oxc* and *oxlT.* Thanks to the construction and validation of specific conservation models for all three genes, we were able to search for oxalotrophy in genomes and metagenomes associated with the human digestive tract, oral cavity, and lungs. We report that oxalotrophy - the capacity to use oxalate as a source of energy - is a rare metabolic trait, mostly confined to the gut, and also find evidence that it can be acquired by horizontal gene transfer. By contrast, the capacity for oxalate degradation is more widespread, and two genes responsible for it (*frc* and *oxc*) are almost always close together in the genome, suggesting selection pressure.

## Introduction

Oxalate — the fully deprotonated form of oxalic acid at physiological pH — occurs in various roles in the metabolism of a wide array of organisms. It can serve, for example, as a source of energy in bacteria (Unden 2013), as a virulence factor in plant pathogenic fungi (Wang and Wang 2020) or as a defense compound in plants (Franceschi 2001). In humans, the association of oxalic acid with disease is of considerable importance. The amount of oxalate in circulation in the human body depends on multiple factors including dietary intake, intestinal absorption, endogenous production, and renal excretion (Baltazar et al. 2023). Metabolic alterations can lead to toxic accumulation of oxalate in body tissues. The kidney is the main excretory organ and a primary target of oxalate toxicity (Rosenstock et al. 2022). Calcium oxalate crystals are commonly found in kidney stones, but they are also found in over two thirds of endemic bladder stones (Leslie et al. 2025).

Humans lack endogenous metabolic pathways involved in the degradation of oxalate. Instead, the degradation of oxalate by bacteria in the gastrointestinal tract is thought to decrease the rates of absorption (Karamad et al. 2022; Allison et al. 1986). In fact, one of the model organisms that allowed the study of oxalate catabolism is *Oxalobacter formigenes*, a bacterium found in the digestive tract of vertebrates (including humans). This organism is the model for *oxalotrophy,* which, in the present work, is defined as the capacity to use oxalate as a source of *energy* through virtual proton pumping (see next paragraph). Other authors may use the term to denote different metabolic pathways involving oxalate, which may or may not be linked to the generation of energy for growth. For example, Sahin (2003) defines the term as the capacity to use oxalate as the *sole* source of carbon and energy, but without restriction to a specific pathway.

Oxalate catabolism in *O. formigenes* has been known for several decades and is relatively well described. *O. formigenes* is able to create a transmembrane electric potential difference by exchanging one oxalate molecule (carrying two negative charges) with one formate molecule (carrying one negative charge) (Ruan et al. (1992) - see figure **1**). Contrary to respiration, there is no electron transport chain. The electric potential difference then drives the synthesis of ATP by an ATP synthase in a process known as a *virtual proton pump* (figure **1**, panel A). In the case of oxalotrophy, the imported oxalate is decarboxylated to formate and carbon dioxide by the successive action of formyl-CoA:oxalate-CoA transferase and oxalyl-CoA decarboxylase. These are encoded by the *frc* and *oxc* genes, respectively. The decarboxylation step is coupled with the transport by the oxalate:formate antiporter, a transmembrane protein that belongs to the major facilitator superfamily, encoded by the *oxlT* gene (figure **1**, panel B). Overall, the balance is similar to that of a dismutation as found in fermentations, with a free energy of-26.2 kJ/mol (figure **1**, panel C).

**Figure 1.**
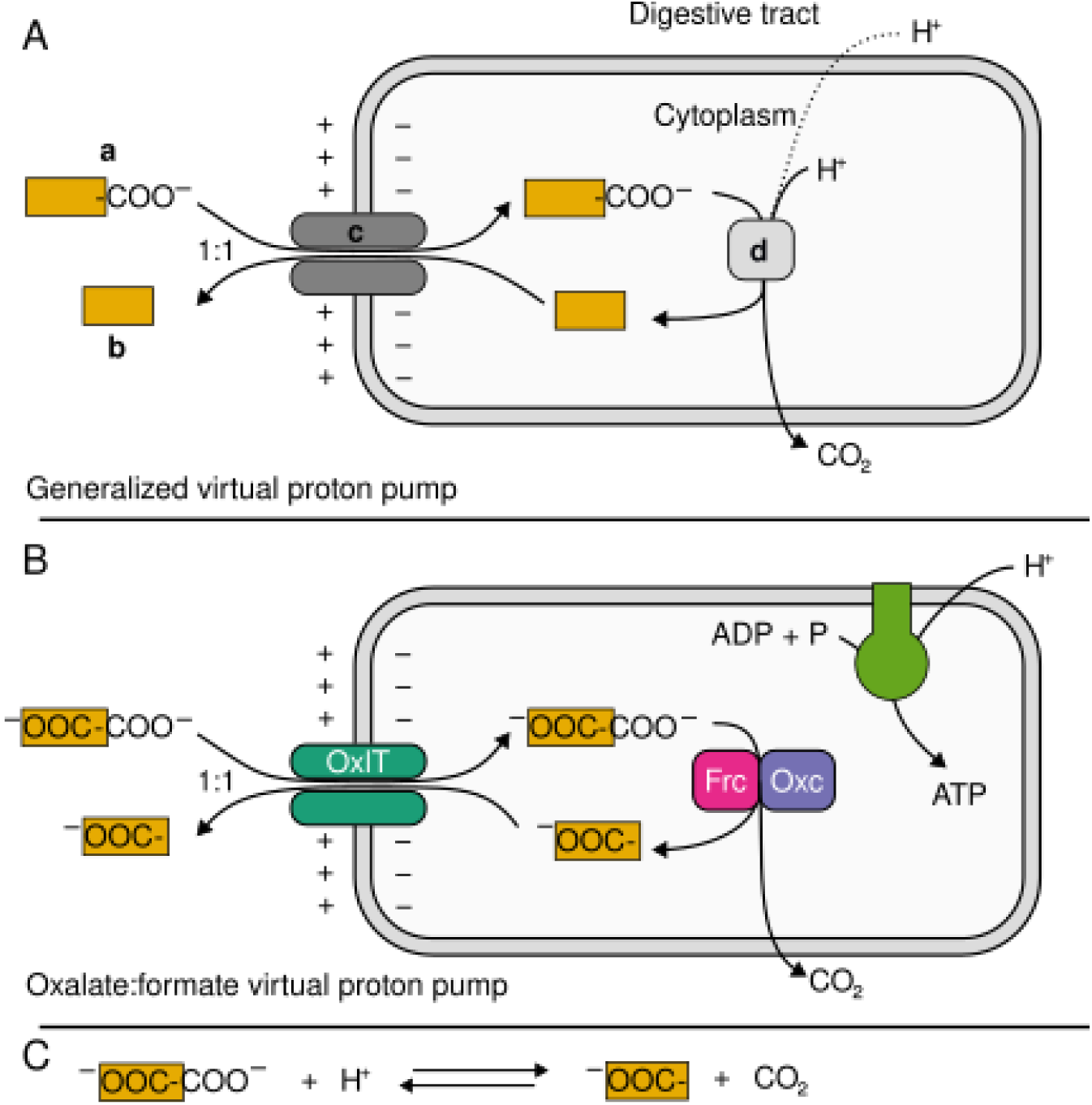
A virtual proton pump (panel A) involves two molecules, one of which (**b**) is the decarboxylated form of the other (**a**). A membrane-bound antiporter (**c**) imports one **a** while it exports one **b**. Since **a** has one more negative charge than **b**, the exchange results in the net accumulation of one positive charge on the exterior of the membrane. Within the cell, **a** is decarboxylated into **b** by one or more cytoplasmic enzymes (**d**). The reaction is driven by the free energy of decarboxylation (Fang et al. 2009).

In the oxalate:formate virtual proton pump discussed in this paper (panel B), **a** and **b** are oxalate and formate, respectively, and the virtual protons contribute to the transmembrane electrical potential dissipated by the cell to synthesize ATP. Other systems exist, such as the acid-resistance response of *E. coli*, in which **a** is L-arginine and **b** is agmatine. In this case, the ejection of protons from the cytoplasm amounts to keeping its pH tolerably high.

The overall reaction for oxalate:formate decarboxylation is shown in panel C.

The prevalence of oxalotrophy among members of the human microbiota has not been extensively investigated. A recent study assessing the frequency of oxalate-degrading pathways in the human gut microbiota showed a widespread detection of the *frc* and *oxc* genes (Liu et al. 2021). However, a search for the simultaneous presence of homologs of *frc*, *oxc*, and *oxlT* within the same genome, which is a necessary condition for an organism to be considered an oxalotroph in the sense used here, has never been made. The aim of the present study was therefore to develop and validate an approach to scan genomes and metagenome-assembled genomes (MAGs) from the human microbiome for candidate oxalotrophs. Scanning for a gene in a genome (or for a protein a proteome) is best performed using profiles (Lüthy et al. 1994) or hidden Markov models (Durbin et al. 1998), although simpler, less powerful methods exist, such as position-specific weight matrices (Gribskov et al. 1987). All of the above are mathematical representations of conserved sequences. To refer to any such representation of sequence conservation, we hereafter use the generic term *model,* for conciseness.

Scanning for a gene or protein is straightforward if an adequate model already exists, for example in a public database such as ProSite (Sigrist et al. 2012) or InterPro (Paysan-Lafosse et al. 2022). However, not all genes and proteins have been modeled. Furthermore, existing models may be either too restricted or too lax, as when a model targets a whole protein family *vs.* a specific member. Accordingly, a secondary objective of the present study was to ensure that adequate models exist for all three genes responsible for oxalotrophy.

## Materials and Methods

### Genomes and MAGs

The source sequences consisted of a combination of genomes and metagenomes of representatives of the human microbiota (table 1). Those included 4,728 representative MAGs obtained from the catalog of human gut microbiome (Almeida et al. 2021), 452 MAGs from the oral cavity (Gurbich et al. 2023); and 18 genomes from the lung (Das et al. 2021). In addition, a set of six genomes from oxalotrophic bacteria was used as a positive control set (F. Palmieri, pers. comm, Table **2**).

**Table 1:** Sources of Primary Data.

| Set | Count | Source |
| --- | --- | --- |
| Lung | 18 | (Das et al. 2021) |
| Oral cavity | 452 | v1.0.1; (Gurbich et al. 2023) |
| Digestive tract | 4,728 | v2.0.1; (Almeida et al. 2021) |
| Total | 5,198 |  |

**Table 2:** Known Oxalotrophs (Positive Control Set)

| Assembly number | Strain | Evidence for oxalate as sole source of energy? | Isolation Source |
| --- | --- | --- | --- |
| GCF_003609605.1 | *Ammoniphilus oxalaticus* RAOx-1 | (Zaitsev et al. 1998) | *Rumex acetosa* L. rhizosphere (Zaitsev et al. 1998) |
| GCF_000020125.1 | *Paraburkholderia phytofirmans* PsJN | (Kost et al. 2014) | *Allium* sp. rhizosphere (Reimer et al. 2022) |
| GCF_004798725.1 | *Cupriavidus necator* H16 | (Cowan et al. 2024) | soil (inferred from type strain N-1) (Reimer et al. 2022) |
| GCF_008807855.1 | *Cupriavidus oxalaticus* T2 | (Yan et al. 2021) | Activated sludge |
| GCF_004768545.1 | *Cupriavidus oxalaticus* X32 | (Khambata and Bhat 1953) | sludge (Xiang et al. 2020) |
| GCF_002128345.1 | *Oxalobacter formigenes* OXCC13 | (Allison et al. 1985) | human feces (Allison et al. 1985) |

### General Gene Prediction and Annotation

To ensure consistency of annotation, all MAGs were subjected to gene prediction and annotation with Prokka (Seemann 2014). Taxonomy assignment was performed with GTDB-Tk (Chaumeil et al. 2022), version 2.1.1.

### Scanning for InterPro Models

The positive and negative controls were scanned for hits of InterPro models with InterProScan (Jones et al. 2014) using the InterProScan website (https://www.ebi.ac.uk/interpro/search/sequence/). The results were downloaded as a JSON file and processed with `jq` (Färber 2023).

### Search for *frc*, *oxc*, and *oxlT* Genes

The *frc*, *oxc*, and *oxlT* genes were searched for with generalized profiles (Lüthy et al. 1994). This was performed at the protein level for increased sensitivity; that is, protein profiles were matched against translations of predicted genes. The profiles were validated by examining phylogenies of profile hits and confirming that all hits above a certain (protein-dependent) threshold formed a clade which contained all and only sequences with the expected annotations among known sequences.

### Pairwise Sequence Comparisons

These were carried out with Needleman and Wunsch’s algorithm (Needleman and Wunsch 1970) as implemented in the `needle` program from the EMBOSS (Rice et al. 2000) package, using default parameters.

### Gene Neighbourhoods

The gene neighbourhoods were plotted with `gggenes` (David 2023), an extension of the `ggplot2` (Hadley 2016) R package, itself part of the `tidyverse` collection (Wickham et al. 2019).

### Phylogenetic Analysis

#### Multiple Alignments

Multiple alignments were computed with MAFFT (Katoh et al. 2005), using default parameters.

#### Phylogenetic Trees

Phylogenetic trees were computed with IQ-TREE (Nguyen et al. 2015). Model selection was performed with ModelFinder (Kalyaanamoorthy et al. 2017). Support values were computed as ultrafast bootstraps and are represented as a percentage out of 1,000 replicates. The trees were rendered as graphics either with the `ggtree` (Yu 2020) package for R (R Core Team 2024) or the Newick Utilities (Junier and Zdobnov 2010).

#### Shared Bipartitions

Bipartitions shared by a pair of trees (a measure of similarity) were counted with the ComparePhylo() function from the APE (Paradis and Schliep 2019) R package.

#### Optimization of the Profile Hit Set for maximal Shared Bipartition Number

The set of profile hits yielding the maximal number of shared bipartitions (See Results and Discussion) was identified using a genetic algorithm implemented in the kofnGA (Wolters 2015) R package.

## Results and Discussion

### Search for a suitable set of models for Frc, Oxc, and OxlT in public databases

A prerequisite for scanning genomes for oxalotrophy is a set of good quality oxalotrophy protein models. Quality, in this case, means in particular the capacity to distinguish between positive controls and their closest relatives, which by definition are very similar at the sequence level, and hence a likely source of false positives. An ideal set of models would also come from the same database, because models reflect the choices of researchers who construct them (such as the programs used to build the models, and the parameters of these programs): models from different sources are therefore more likely to embody different methodologies, and are thus inherently less comparable than models from the same source.

To gauge the accuracy of models from public databases, we constructed positive and negative control sets, for each of Frc, Oxc, and OxlT - the gene products of *frc*, *oxc*, and *oxlT*. For this, positive and negative controls were identified based on the Prokka annotations of all genomes and MAGs. Controls were then subjected to (i) an InterPro scan, which returned all InterPro models with at least one match in one of the control sets, and (ii) a construction of a phylogeny. To keep the phylogeny amenable to visual inspection, we limited the sets to 30 positive and 70 negative controls, drawn at random. Figure **2** shows a subset of the results for Oxc; the complete figures are available in the supplementary data as Supplementary Figures 1, 2 and 3. More than 80 models had at least one match among the controls. Of these, only one (namely, PTHR43710:SF2, from the Panther database(Mi et al. 2021)) matched all positive controls and no negative controls - all others either missed at least one positive or matched at least one negative control. Moreover, PTHR43710:SF2 is actually not a model for oxalyl-CoA decarboxylase - instead, it targets 2-hydroxyacyl-CoA lyase, which is a different enzyme (Thomas et al. 2022). In other words, none of the models for oxalyl-CoA decarboxylase had 100% precision and recall, and the only model that does achieve this performance is for a different target. Therefore, it is not obvious that PTHR43710:SF2, despite its excellent precision and recall on this particular set, is a good choice as a model for Oxc. Similar results were obtained for OxlT and Frc (see supplementary figures **1** and **3**). It was unclear, therefore, that there existed models for all three proteins corresponding to oxalotrophy genes, of whose accuracy we could be confident. It seemed even less clear that such a set could come from the same database. Accordingly, we built new, in-house models.

**Figure 2.**
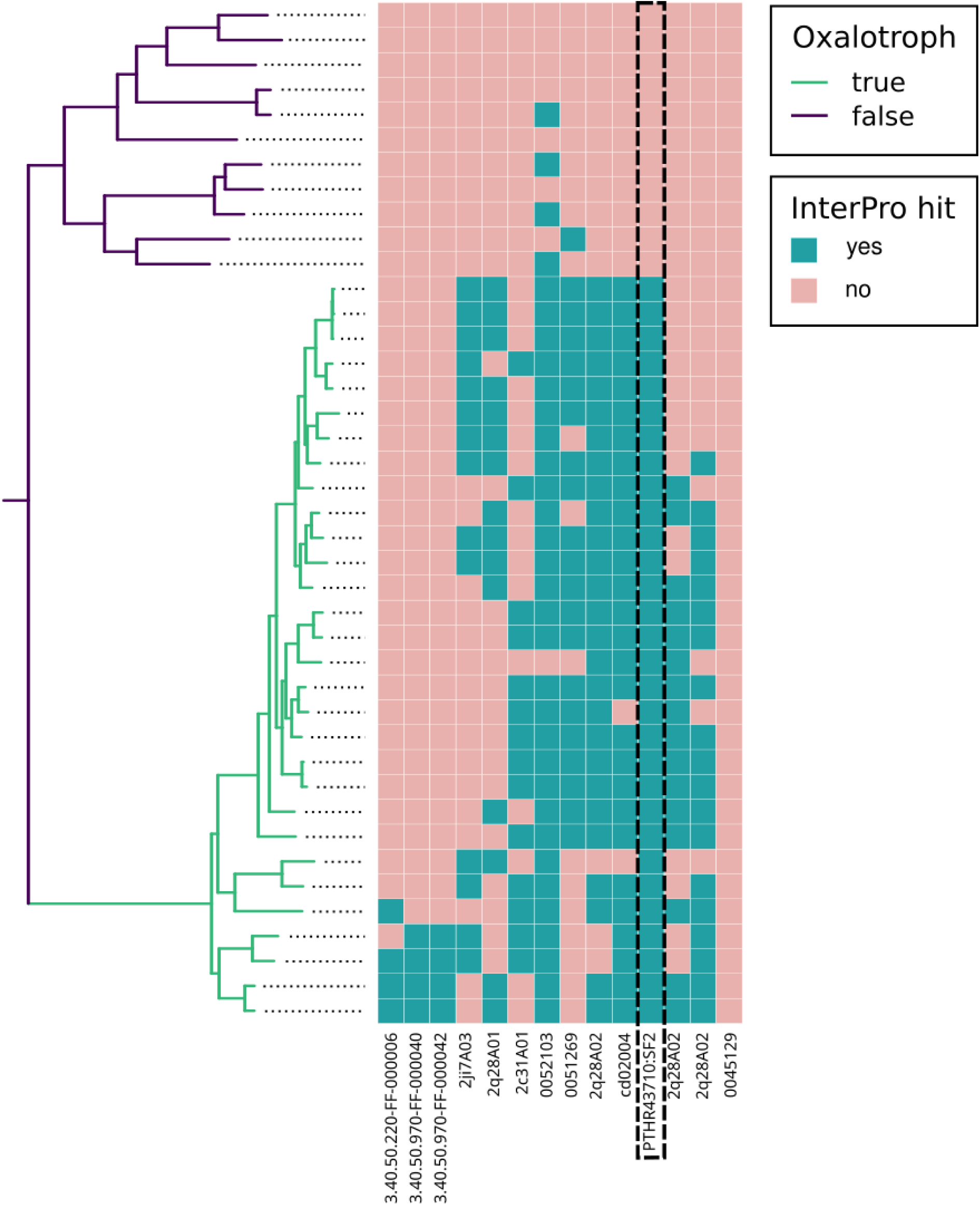
Hits of InterPro models of Oxc on the sequences in the control sets. Each column in the heatmap pertains to one model, and each line corresponds to a protein in the tree on the left. A cell in the heatmap is coloured cyan if the corresponding model has at least one hit in the corresponding protein. The clade of true positives in the tree is coloured green, the rest is in purple. To allow the figure to fit on a page, only a fraction of the negative controls, and only some models, are shown.

Model PTHR43710:SF2 (and only this model — dashed box) has precision and recall both 1.0 (all true positives and negatives correctly identified), however it does not target oxalate-CoA decarboxylase, but 2-hydroxyacyl-CoA lyase. The implication is that none of the models available in public databases is entirely suitable for oxalate-CoA decarboxylase.

### Development of new Models for Frc, Oxc, and OxlT

For the new models, we chose the profile method, as implemented in the Pftools (Schuepbach et al. 2013). Aims for the new profiles were that (i) all and only positive hits correspond with Prokka annotations, and (ii) all and only positive hits form a clade in a phylogeny of homologs. The profile method allows the use of two score thresholds (see Schuepbach et al. (2013) and references therein for details): a *lower* one, intended to recognize a wide set of homologous sequences, and a *higher* one, used to recognize the target set itself. During the validation process, we constructed phylogenies of all sequences reported at the lower threshold, thus deliberately including not only our genes of interest, but also their nearest relatives, which count as false positives. We then verified that the sequences with hits above the higher threshold formed a clade among the larger set (see Supplementary figures **4-6**). Were this not the case, there would be reason to doubt that any sequence was a member of the target set, even with a profile score above that threshold.

### Screening for Frc, Oxc, and OxlT

The new profiles were searched for in genomes and MAGs from the human digestive tract, lung, and oral cavity, as well as the controls.

#### Occurrence Per Phylum

Of the 5198 genomes and MAGs screened (see table **1**), 10 contained all three genes required for oxalotrophy, and are therefore potential oxalotrophs, while 125 lacked *oxlT* but did contain *frc* and *oxc* (table **3**). Of the 10 potential oxalotrophs, 2 were strains of *O. formigenes*, while another 3 were *Oxalobacter* but from undetermined species. This makes *Oxalobacter* strongly overrepresented among the potential oxalotrophs. The 6 members of the control set (see table **3**) also contained all three genes, as expected. In other words, among the members of the human microbiota, species with the complete oxalotrophy gene set were very rare (10 / 5198 ≈ 0.19 %), while those capable of decarboxylating oxalate slightly less so 125 / 5918 ≈ 2.5 %). All the potential oxalotrophs were found among the MAGs from the gut dataset. The potential oxalotrophs belonged to phyla Firmicutes or Proteobacteria, while the 125 species possessing *frc* and *oxc* belonged to Actinobacteriota, Bacteroidota, Desulfobacterota I, Firmicutes, and Proteobacteria (table **4**).

**Table 3.** Species/MAGs with hits of all profiles. . These fall into two groups: (i) genomes from the control set (white background) - all six were recovered, as expected; (ii) MAGS (grey background) - all originate in the gut dataset, but the isolation source is not always the digestive tract (see text).

| Accession Number | Species | Phylum | Isolate Source |
| --- | --- | --- | --- |
| GCF_003609605.1 | *Ammoniphilus oxalaticus* | Firmicutes | rhizosphere |
| MGYG000001265 | *Methylobacterium*<br>sp002778835 | Proteobacteria | unknown (genus reported from soil, water); possible contaminant |
| MGYG000002310 | *Microvirga massiliensis* | Proteobacteria | human feces |
| MGYG000003734 | *Afipia broomeae* | Proteobacteria | human sputum |
| MGYG000003137 | *Bradyrhizobium denitrificans* | Proteobacteria | freshwater lake |
| MGYG000003733 | *Bradyrhizobium*<br>sp003020075 | Proteobacteria | human gut |
| GCF_004798725.1 | *Cupriavidus necator* H16 | Proteobacteria | soil |
| GCF_008807855.1 | *Cupriavidus oxalaticus*<br>T2 | Proteobacteria | sludge |
| GCF_004768545.1 | *Cupriavidus oxalaticus*<br>X32 | Proteobacteria | sludge |
| MGYG000003451 | *Oxalobacter* sp. | Proteobacteria | unknown |
| GCF_002128345.1 | *Oxalobacter formigenes*<br>OXCC13 | Proteobacteria | human gut |
| MGYG000001331 | *Oxalobacter formigenes* | Proteobacteria | human gut |
| MGYG000002505 | *Oxalobacter formigenes*<br>B | Proteobacteria | human gut |
| MGYG000004063 | *Oxalobacter*<br>sp900760095 | Proteobacteria | unknown |
| MGYG000002703 | *Oxalobacter*<br>sp9052202055 | Proteobacteria | unknown |
| GCF_000020125.1 | *Paraburkholderia phytofirmans* PsJN | Proteobacteria | rhizosphere |

**Table 4.** Number of genomes/MAGs containing *frc*, *oxc*, and *oxlT.* Column 1: phylum; column 2: number of species per phylum among the genomes/MAGs containing both *frc* and *oxc*; column 3: number of species containing *oxlT* as well.

| Phylum | # Genomes/MAGs containing *frc* and *oxc* | # Genomes/MAGs containing *frc*, *oxc* and *oxlT* |
| --- | --- | --- |
| Actinobacteriota | 8 | 0 |
| Bacteroidota | 75 | 0 |
| Desulfobacterota I | 2 | 0 |
| Firmicutes | 14 | 1 |
| Proteobacteria | 26 | 15 |
| Total | 125 | 16 |

#### Ecological Origin

The overrepresentation of *O. formigenes* or other species related to *Oxalobacter* is in agreement with previous studies indicating that members of this genus dominate the function of oxalate degradation in the human gut. The presence and expression of oxalate-degrading enzymes in metagenomes and metatranscriptomes of diverse human cohorts (Liu et al. 2021). This also confirms previous studies that had shown a correlation between the absence of this species and a deregulation of oxalate homeostasis (Sidhu et al. 1999). Moreover, the detection of other potential novel oxalotrophs is noteworthy. One MAG with hits by the three profiles was classified as *Afipia broomeae*, a species that has been previously isolated from sputum, and may therefore be associated with the oral cavity. Representatives of the genus *Afipia* have been shown to grow on oxalate as sole source of carbon, but also to degrade calcium oxalate *in vitro.* The same has been reported for representatives of the genus *Bradyrhizobium* (Cowan et al. 2024) which could also correspond to oxalotrophs naturally occurring in the oral cavity. In contrast, MAGs related to organisms such as *Bradyrhizobium denitrificans* or the two *Methylobacterium*, which have not been reported from the gut microbiome (Table **3**), could be the result of contamination (*Methylobacterium* is a known contaminant of genomic datasets (Salter et al. 2014)), but also to the presence of a transient microbiota related to food items. For instance, *Methylobacterium* are among methylotrophic oxalotrophs that are known as plant colonizers and for which oxalate represents an important carbon source in plant-associated habitats (Schneider et al. 2012).

#### Gene Copy Number

In the 16 species possessing all three oxalotrophy genes, *oxc* was always single-copy while *oxlT* was always present in at least two copies; *frc* being more frequently multi-copy than *oxc* but less so than *oxlT* (supplementary table **2**). In the 125 species that contain *frc* and *oxc*, the minimum and median are 1 copy for both genes, while the maxima are 2 and 4 copies, respectively (supplementary table **3**). This finding is noteworthy as the *frc* has been used in the past as a molecular marker not only to characterize, but also to quantify oxalotrophs (Khammar et al. 2009). However, the results obtained here suggest that *oxc* would be a more adequate marker as it would not result in the over-estimation of oxalotrophic bacteria being a single copy gene.

#### Genomic organisation of *frc*, *oxc*, and *oxlT*

In the genomes/MAGs that contain both *oxc* and *frc*, these two genes were found to be adjacent in all Firmicutes, almost all Bacteroidota, Desulfobacteriota I, and the majority of Proteobacteria. Only in Actinobacteria are they always separated, though by at most fourteen genes and usually fewer (Table **7**). The two genes lie predominantly in the same orientation (though not necessarily the same order — see below), suggesting that they may be part of an operon, as shown in the past for some oxalotrophic models such as *Lactobacillus acidophilus* (Azcarate-Peril et al. 2006) or *Cupriavidus oxalaticus* (Palmieri et al. 2022). Here as well the Actinobacteriota are an exception: the two genes lie in opposite orientations. Finally, the order in which *frc* and *oxc* appear on the genome varies *across* Phyla but is relatively stable *within* them, with the exception of Proteobacteria (Table **7**) which are divided into two clades, one with the *frc* → *oxc* organisation (relative to the direction of transcription), and the other with *oxc* → *frc*, or with *oxc* alone (in some *Oxalobacter*) — see Supplementary Figure 7. The latter of these clades contains exactly those Proteobacteria whose genomes also contain *oxlT*, and apart from Proteobacteria includes the Firmicute *Ammoniphilus oxalaticus*. The overall conservation of the genomic organisation of *frc* and *oxc* across phyla as distant as Firmicutes and Proteobacteria suggests a selection pressure that keeps these two genes in close proximity possibly as an operon (but see the case of *O. formigenes* below) In genomes that also contain *oxlT*, a copy of this gene is usually found in the vicinity of *frc* and *oxc*, and usually in the same orientation, which as before suggests that they are part of an operon. Again, *Oxalobacter* is an exception: *oxlT* may be found in the vicinity of either *frc* or *oxc* in *O. formigenes*, but in other *Oxalobacter* MAGs the genes are not in the vicinity of one another. This may be linked to the fact that at least some *Oxalobacter* are *obligate* oxalotrophs: the oxalotrophy genes might be constitutively expressed, and might therefore not need to be organized in an operon since expression would not be a response to a signal. Finally, other genes closely related to *oxlT* may be found in the vicinity of *frc* and *oxc* as well, such as the L-lactate transporter and other members of the major facilitator superfamily (not shown).

**Table 7.** Genomic organisation of *frc* and *oxc*. Column 1: phylum; Column 2: genomic organisation of *frc* and *oxc*, in 5’-3’ orientation. Column 3: number of MAGs in which the structure was found. The two genes are predominantly adjacent, or else they are separated by at most 14 genes - only in 6 Proteobacteria and 8 Bacteroidota are they more widely separated (represented by *oxc* alone in the table cell). In Actinobacteria, *oxc* and *frc* have opposite orientations (shown as *<u>frc</u>* (underlined)).

| Phylum | Genomic structure | Count |
| --- | --- | --- |
| Actinobacteria | *oxc* → 2xN → <u>*frc*</u> | 4 |
|  | *oxc* → N → <u>*frc*</u> | 3 |
|  | *oxc* → 14xN → <u>*frc*</u> | 1 |
| Bacteroidota | *frc* → *oxc* | 67 |
| Bacteroidota | *oxc* | 8 |
| Desulfobacteriota I | *frc* → *oxc* | 2 |
| Firmicutes | *oxc* → *frc* | 13 |
| Firmicutes | *frc* → *oxc* | 1 |
| Proteobacteria | *oxc* → *frc* | 9 |
| Proteobacteria | *frc* → *oxc* | 7 |
| Proteobacteria | *frc* → *oxc* → N → *frc* | 6 |
| Proteobacteria | *oxc* | 6 |

### Phylogeny of Frc, Oxc, and OxlT among species that contain all three proteins

Phylogenies were built from all copies of the Frc, Oxc and OxlT sequences from the sixteen species that contain at least one copy of each. In each case, the phylogeny was rooted on the true positive clade’s sister clade (see supplementary figures **4-6**). All these species belong to phylum Proteobacteria, except *A. oxalaticus*, which is a Firmicute. This is not always reflected in the phylogenies, however: the four copies of Frc from *A. oxalaticus*, for example, are most closely related to their homologs from *Oxalobacter* (Figure **3** — indeed, they are the closest relatives of *Oxalobacter*’s Frc, notwithstanding the fact that the rest of the tree comprises only Proteobacteria. For Oxc, the phylogeny does place *Ammoniphilus* as a sister to all Proteobacteria, as would be expected (Figure **4**); however, a phylogeny with more Proteobacterial species again places *A. oxalaticus’* Oxc within the Proteobacteria (see Supplementary Figure 7). As for OxlT, the situation is similar to that of Frc (Supplementary Figure 8). There is therefore indication that all of the oxalotrophy genes of *A. oxalaticus* are of Proteobacterial origin, and were acquired by horizontal gene transfer, possibly from *Oxalobacter* or a closely-related genus. This aligns with a recent study (Sonke and Trembath-Reichert 2023) that reported phylogenies with subtrees grouping by environment rather than taxonomy. This suggests that while oxalotrophy is generally a vertically-inherited trait, gene transfer remains a possible mechanism in these environments.

**Figure 3:**
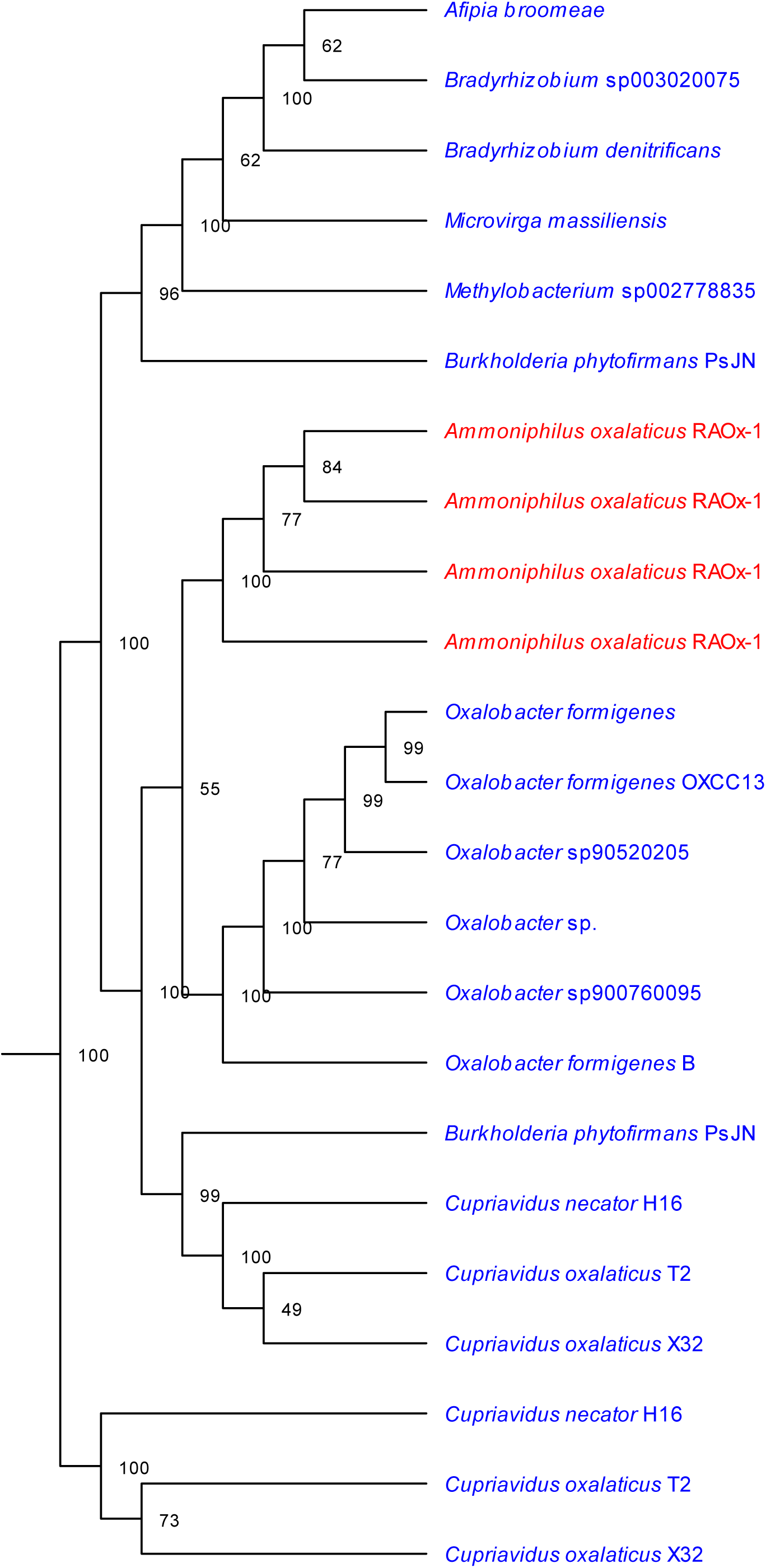
Phylogeny of Frc among the 16 species that contain *oxc*, *frc* and *oxlT*. The tree has more than 16 leaves because most species have more than one copy of *frc*. Blue: Proteobacteria, red: Firmicutes. *A. oxalaticus* (red) has four copies of *frc*, all most closely related to their homologs from *Oxalobacter* and themselves the closest relatives of *Oxalobacter* spp.’s *frc*. This suggests that they were acquitted by *Ammoniphilus* from a Proteobacterial donor. See also figure 3 and supplementary figure 8, as well as supplementary figure 7.

**Figure 4:**
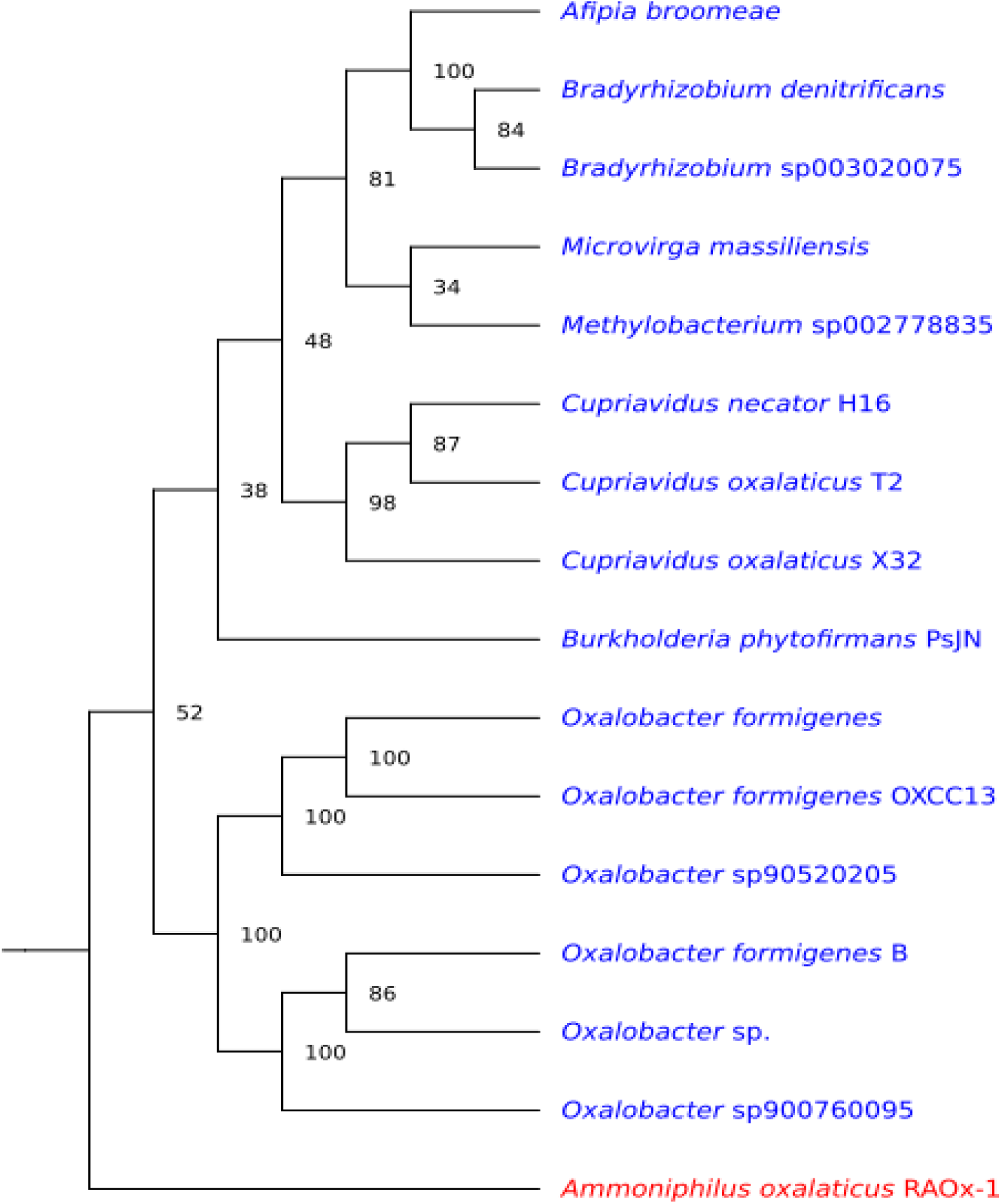
Phylogeny of Oxc among the 16 species that contain *oxc*, *frc* and *oxlT*. The tree has exactly 16 leaves since *oxc* is single-copy in all species. Blue: Proteobacteria, red: Firmicutes. *A. oxalaticus* (red) is outside of the Proteobacterial clade (but see figure 4 and supplementary figure 8, as well as supplementary figure 7).

### Similarity of the Frc, Oxc, and OxlT phylogenies

As seen above, there is indication that the three genes for oxalotrophy may undergo horizontal gene transfer. If this is the case, some degree of coevolution between the genes should be expected. The phylogenies of two coevolving genes are expected to be more similar to each other than are the phylogenies of two genes (from the same species set) that do not undergo coevolution. The similarity of the phylogenies of each of Frc, Oxc, and OxlT to the other two was therefore measured by counting the number of bipartitions shared by each pair of phylogenies. Shared bipartitions are only defined between trees built on exactly the same species, so to enable this comparison, each phylogeny had to contain exactly one sequence from each of the suitable species. This was straightforward for *oxc*, which is single-copy in all species, but not for *frc* and *oxlT*, which are generally multi-copy, requiring one copy (from each) to be picked. Drawing a copy at random would have been suboptimal because phylogenies computed with different copies of a gene (for the same species) might have different topologies - and thus different bipartitions shared with other phylogenies. It was therefore necessary to identify the topologies yielding the highest number of shared bipartitions. This was carried out with a genetic algorithm (see Methods). To assess the significance of these bipartition counts, the optimisation step was performed again, but after shuffling the species in the trees: any shared bipartitions identified between two shuffled trees can only be the result of chance and therefore provide a comparison point to the number of shared bipartitions counted among actual trees. Pairs of real trees shared from five to seven bipartitions, while pairs of shuffled trees shared no bipartitions (table **9**).

**Table 9.** Number of shared bipartitions after genetic algorithm optimisation. Randomizing the tree by shuffling the species caused the total loss of shared bipartitions. This suggests that the trees are more similar than expected by chance, in turn suggesting coevolution due to selective pressure.

| Protein tree pair | Maximal Number of Shared Bipartitions |
| --- | --- |
| Oxc, Frc | 7 |
| Oxc, OxlT | 5 |
| Frc, OxlT | 5 |
| (shuffled) | 0 |

### Phylogeny of Oxc among the Species with *frc* and *oxc*

The phylogeny of the translated decarboxylase gene from the 125 species that contain both *frc* and *oxc* shows a near-perfect separation of the five phyla to which the 125 species belong (see supplementary figure **7**). The exceptions include the two Desulfobacterota I, which form a clade nested within the Bacteroidota, and one Firmicute, *A. oxalaticus*, which is found within the Proteobacteria. Pairwise sequence comparison of *A. oxalaticus’* Frc with its homologs from all Proteobacteria and all other Firmicutes in the set showed that the *Ammoniphilus* sequence is indeed closer to all Proteobacterial sequences than it is to any (non-*Ammoniphilus*) Firmicute sequence. This suggests that the *Ammoniphilus* sequence is of Proteobacterial origin and was acquired by horizontal gene transfer. This is further supported by the ordering of the *oxc* and *frc* genes in *A. oxalaticus* (namely *frc* - *oxc*): while it is found in nearly half of the Proteobacterial sequences, it is not found in any other Firmicute in our dataset.

### Oxalate metabolism in absence of *oxlT*

Most of the species that contain *frc* and *oxc* do not contain *oxlT*. They therefore most likely do not implement a virtual proton pump, and probably do not use oxalate as a source of energy.

Oxalate might serve as a source of carbon (for example by the conversion of oxalyl-CoA to glyoxylate by glyoxylate dehydrogenase [EC 1.2.1.17] as part of the oxalate degradation III pathway); it may also undergo degradation as a response to acidity, as in the acid tolerance response (ATR) found in *E. coli* (Fontenot et al. 2013). In this case it seems not to be used as a resource by the cell.

### Other functions of Frc

Apart from oxalate, formyl-CoA:oxalate-CoA transferase can transfer formyl-CoA to succinate (Baetz and Allison 1990), suggesting the possibility of using this substrate in the acid tolerance response. However, these results were obtained in the laboratory, and it is not known whether this reaction occurs in the natural environment.

### Origin of the Virtual Proton Pump

Since *frc* and *oxc* are found in many phyla of Bacteria, while *oxlT* seems restricted to Proteobacteria (or, in the case of *Ammoniphilus*, was acquired from them), a parsimonious scenario would be an original, widespread capacity for oxalate decarboxylation via a CoA intermediate, followed by the development of the virtual proton pump in Proteobacteria.

## Conclusions

The ability to use oxalate as a source of energy by virtual proton pumping, as inferred by the presence of *frc*, *oxc* and *oxlT* in the genome, is rare among the bacteria associated with humans, and was not found outside the digestive tract. One candidate (*A. broomeae*) was isolated from the oral cavity; the others are enteric. A larger, but still small, fraction of the genomes studied lack *oxlT* but contain the other two genes. In these organisms, the decarboxylation of oxalate likely has a different function from the biosynthesis of ATP, or else *frc* might accept succinate (rather than oxalate) as a substrate.

When present, *frc* and *oxc* almost always occur next to each other, and at most a dozen genes apart. This suggests at least some selection pressure to maintain the two genes close together on the genome, and possibly to express them together, either constitutively or in response to the same signal. This suggests that a pathway involving the decarboxylation of oxalate with coenzyme A as a cofactor appeared first, and was only later co-opted to function as a virtual proton pump. The order in which *frc* and *oxc* appear on the genome varies across Phyla, but is broadly conserved within them (with the exception of Proteobacteria). In species that also contain *oxlT,* it is usually found in the vicinity of the other two genes. A noteworthy exception to this is the obligate oxalotroph *O. formigenes*, in which the three oxalotrophy genes are not generally found in the vicinity of one another, confirming that obligate oxalotrophy does not imply the presence of an oxalotrophy operon.

The capacity for oxalotrophy may be horizontally transferable from one genome to an other, even distantly related one: this is suggested by the fact that the *frc*, *oxc*, and *oxlT* genes exhibit signs of coevolution, and by the clearly Proteobacterial origins of the *frc*, *oxc*, and *oxlT* orthologs of *A. oxalaticus*, itself a Firmicute.

The simultaneous phylogenomic analysis of *frc*, *oxc*, and *oxlT* with specific profiles was instrumental in reaching the above conclusions. Firstly, the genomic organisation of the genes, including the nearly-universal adjacency of *ocx* and *frc*, would not have been elucidated had the genes been studied in isolation. Secondly, and more importantly, this work would not have been possible if we had selected models from the databases, because these models typically do not target exactly our genes of interest, but either a larger or a narrower scope instead - such as whole protein families.

## Supporting information

Tar archive containing supplementary files.

## Acknowledgements

This research was funded by the Swiss National Science Foundation through the BRIDGE Discovery program under the grant agreement 40B2-0_194701 (P.J., M.P., A.K.) and the Gebert Rüf Stiftung under the grant agreement GRS-064/18 (P.J). We thank Sudip Das and Eric Bernasconi for early access to lung microbiome assemblies. All heavy-duty computing was performed on the University of Lausanne’s HPC cluster Curnagl.

