## Supplementary material for "Prevalence of Oxalotrophy in the Human Microbiome": Tar archive containing supplementary files.: Genomics and Metagenomics of Oxalotrophy_bioRXiv_ST1.docx

**Supplementary Table 1**: Oxalotrophic bacteria of interest

| **code** | **Species** | **oxalotroph** | **genome_ID*** |
| --- | --- | --- | --- |
| Abro | *Afipia broomeae* | yes | MGYG000003734 |
| Aoxa | *Ammoniphilus oxalaticus* RAOx-1 | yes | GCF_003609605 |
| Blac | *Bifidobacterium lactis* DSM-10140 | yes | GCA_000022965 |
| Bden | *Bradyrhizobium denitrificans* | yes | MGYG000003137 |
| Bsp1 | *Bradyrhizobium* sp003020075 | yes | MGYG000003733 |
| Bphy | *Burkholderia phytofirmans* PsJN | yes | GCF_000020125 |
| Csp1 | *CAG-1427* sp905215035 | yes | MGYG000004724 |
| Caer | *Collinsella aerofaciens_*H | yes | MGYG000004692 |
| Cnec | *Cupriavidus necator* H16 | yes | GCF_004798725 |
| Cox1 | *Cupriavidus oxalaticus* T2 | yes | GCF_008807855 |
| Cox2 | *Cupriavidus oxalaticus* X32 | yes | GCF_004768545 |
| Ecol | *Escherichia coli* | yes | MGYG000002506 |
| Eco2 | *Escherichia coli* K-12 | no | GCF_009832885 |
| Halv | *Hafnia alvei* | yes | MGYG000002508 |
| Laci | *Lactobacillus acidophilus* ATCC-4796 | yes | GCA_000159715 |
| Lamy | *Lactobacillus amylovorus* | yes | MGYG000000161 |
| Lsp1 | *Limosilactobacillus* sp012843675 | yes | MGYG000004861 |
| Msp1 | *Methylobacterium* sp002778835 | yes | MGYG000001265 |
| Mmas | *Microvirga massiliensis* | yes | MGYG000002310 |
| Ofo1 | *Oxalobacter formigenes* | yes | MGYG000001331 |
| Ofo2 | *Oxalobacter formigenes OXCC13* | yes | GCF_002128345 |
| Ofo3 | *Oxalobacter formigenes_B* | yes | MGYG000002505 |
| Osp1 | *Oxalobacter sp.* | yes | MGYG000003451 |
| Osp2 | *Oxalobacter sp900760095* | yes | MGYG000004063 |
| Osp3 | *Oxalobacter sp905202055* | yes | MGYG000002703 |
| Pret | *Providencia rettgeri DSM-1131* | yes | GCF_000158055 |
| Pput | *Pseudomomas putida KT2440* | no | GCF_000007565 |

*: MAGs start with MYG
