## Supplementary material for "Prevalence of Oxalotrophy in the Human Microbiome": Tar archive containing supplementary files.: Genomics and Metagenomics of Oxalotrophy_bioRXiv_ST2.docx

**Supplementary Table 2**: **minimum, median, and maximum copy number** of *oxc*, *frc* and *oxlT* among the 16 genomes/MAGs that contain all three. The *oxc* gene is single copy in all genomes, *oxlT* is always present in at least two copies, no gene was found in more than 4 copies.

| **Gene** | **minimum** | **median** | **maximum** |
| --- | --- | --- | --- |
| *oxc* | 1 | 1 | 1 |
| *frc* | 1 | 1 | 4 |
| *oxlT* | 2 | 2 | 4 |
