## Supplementary material for "Prevalence of Oxalotrophy in the Human Microbiome": Tar archive containing supplementary files.: Genomics and Metagenomics of Oxalotrophy_bioRXiv_ST3.docx

#### **Supplementary** **Table 3:** **minimum, median, and maximum copy number** of *oxc* and *frc* among the 125 genomes/MAGs that contain both. The *oxc* gene is single-copy in almost all genomes/MAGs, and reaches a maximum of two copies (in two strains of *Providencia rettgeri*, namely D and DSM-1131); *frc* is also predominantly single-copy, and reaches a maximum of four copies (in *Ammoniphilus oxalaticus*).

| **Gene** | **minimum** | **median** | **maximum** |
| --- | --- | --- | --- |
| *oxc* | 1 | 1 | 2 |
| *frc* | 1 | 1 | 4 |
