## Supplementary figures and images for "Prevalence of Oxalotrophy in the Human Microbiome"

### Abro.pdf

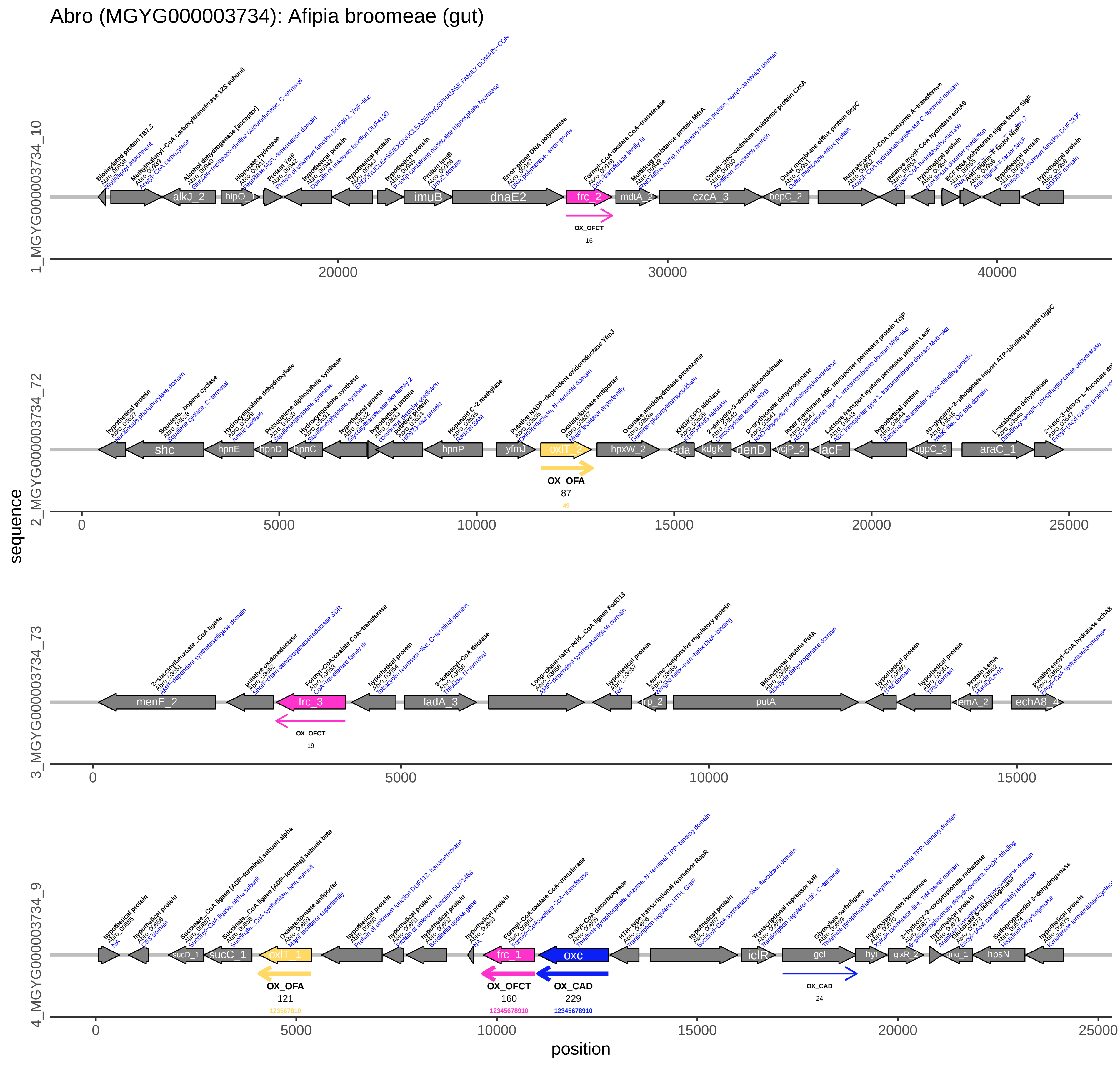

### Aoxa.pdf

#### Aoxa (GCF_003609605): Ammoniphilus oxalaticus RAOx−1 (fabio)

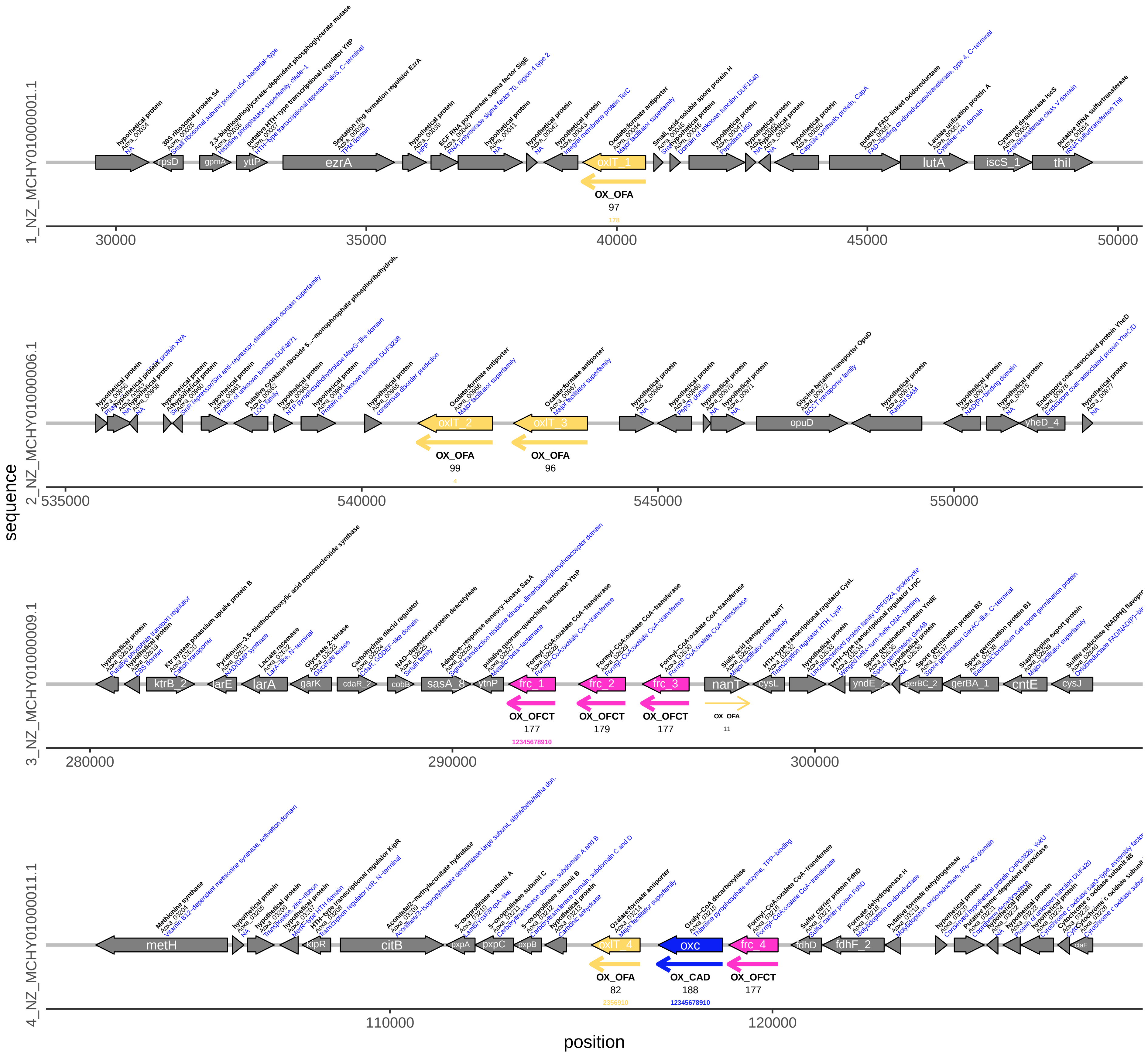

### Bden.pdf

#### Bden (MGYG000003137): Bradyrhizobium denitrificans (gut)

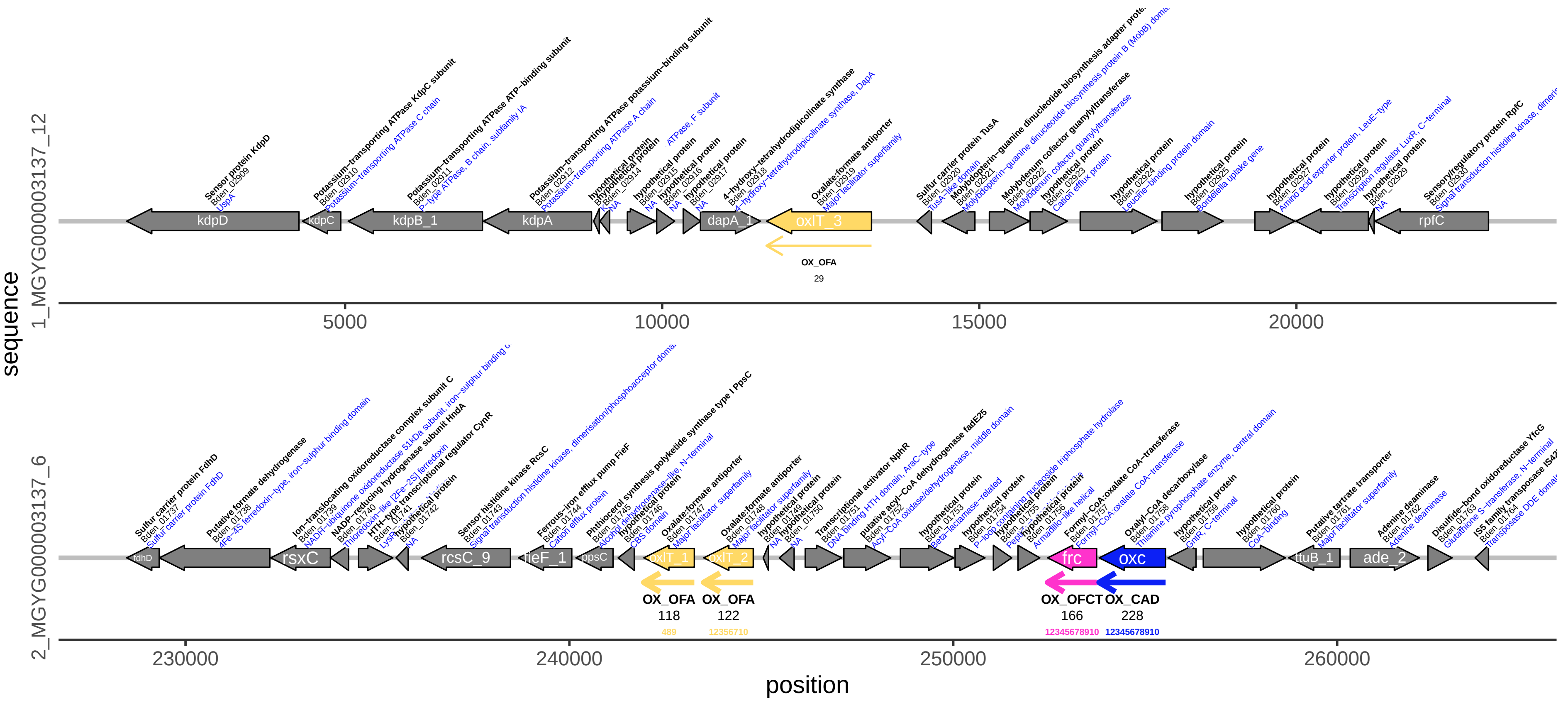

### Bphy.pdf

#### Bphy (GCF_000020125): Burkholderia phytofirmans PsJN (fabio)

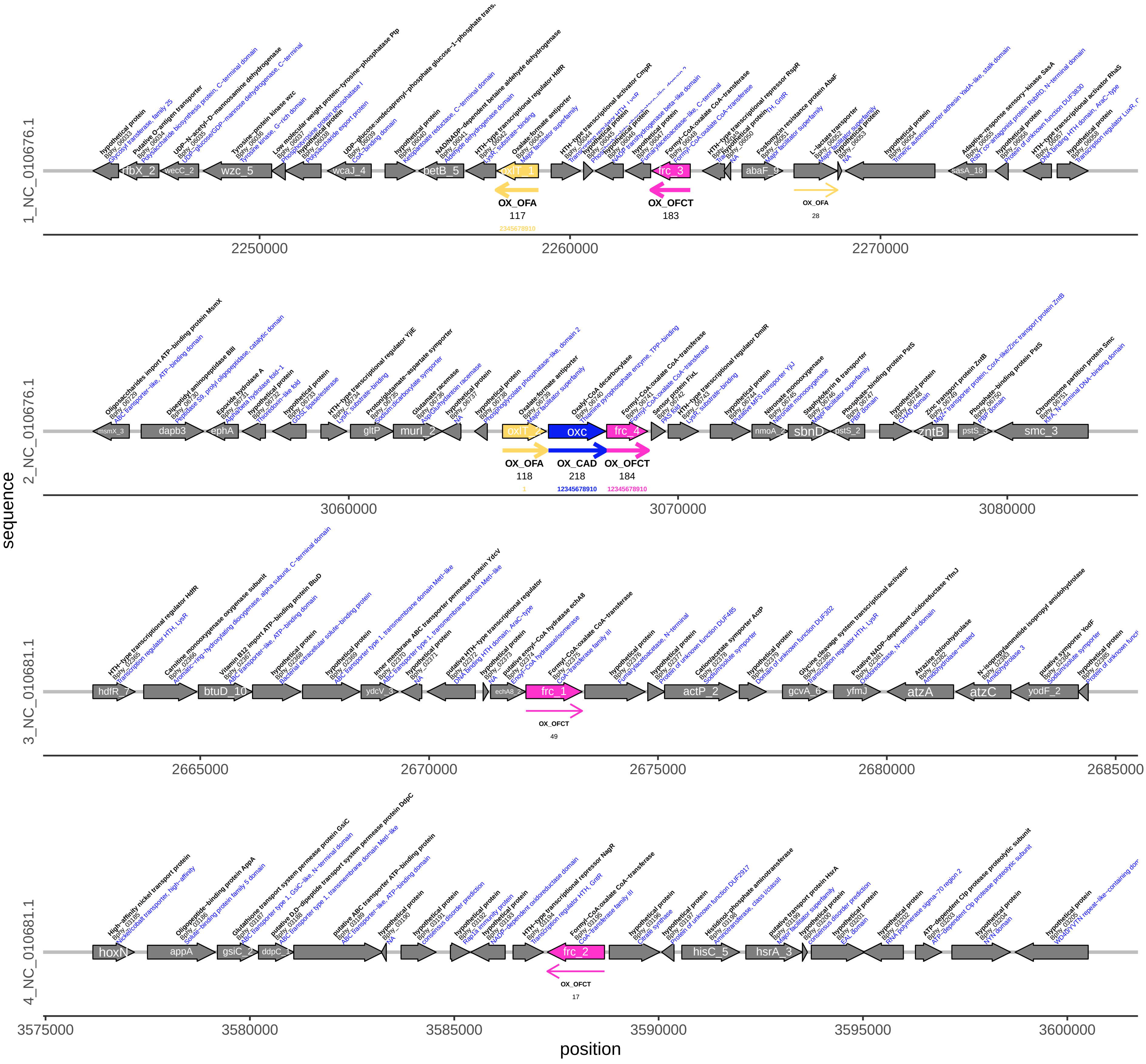

### Bsp1.pdf

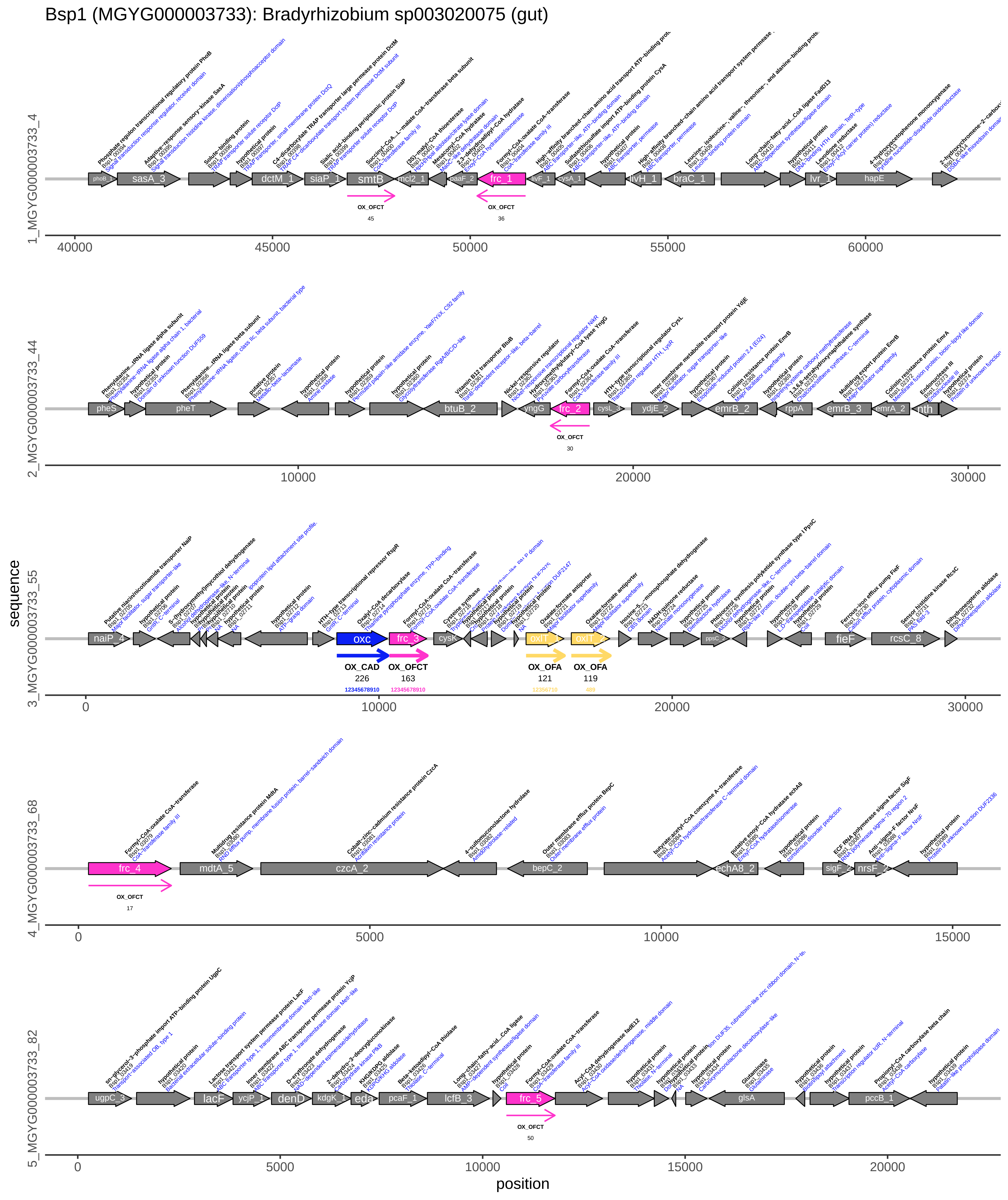

### Cnec.pdf

Cnec (GCF_004798725): Cupriavidus necator H16 (fabio)

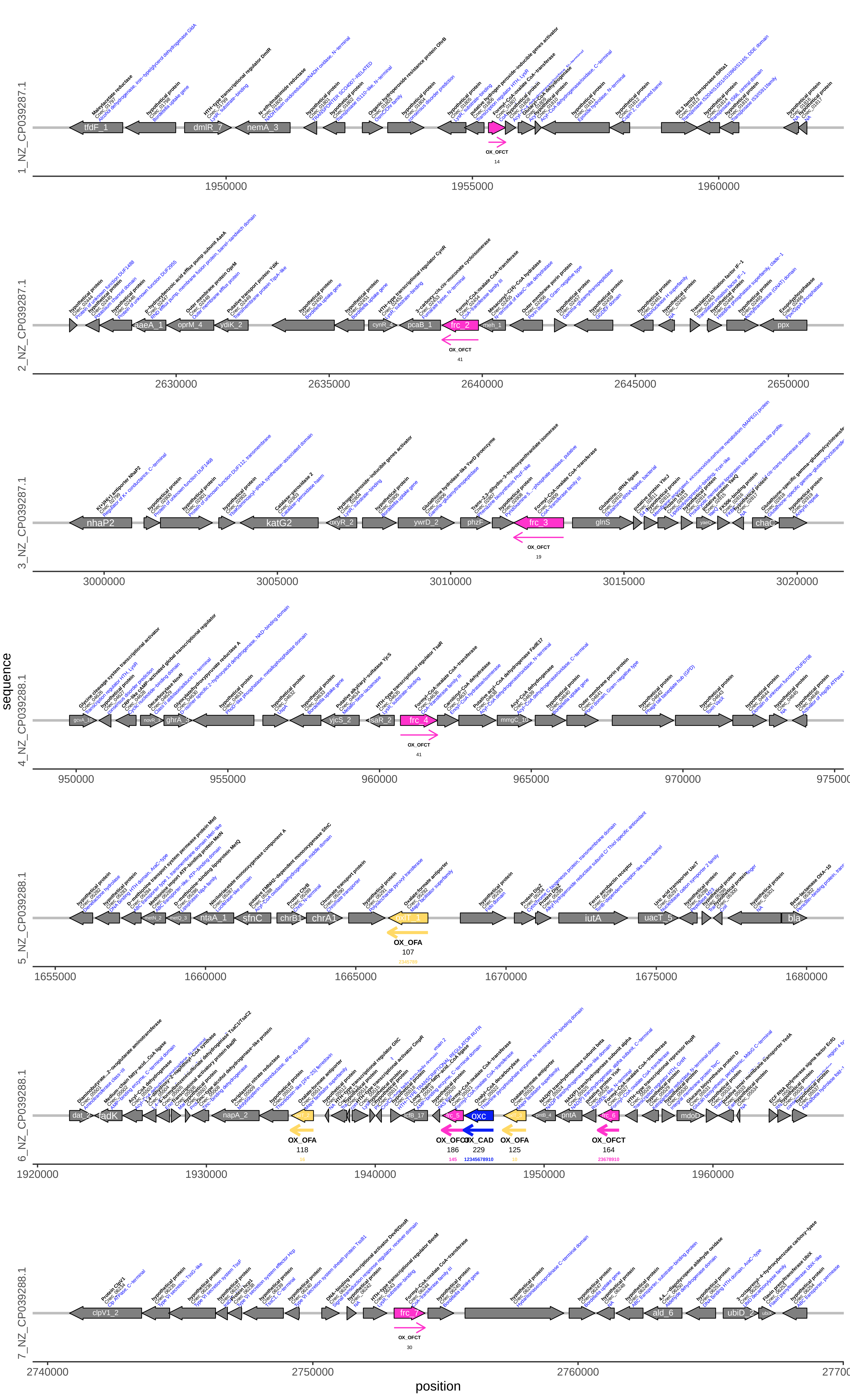

### Cox1.pdf

Cox1 (GCF_008807855): Cupriavidus oxalaticus T2 (fabio)

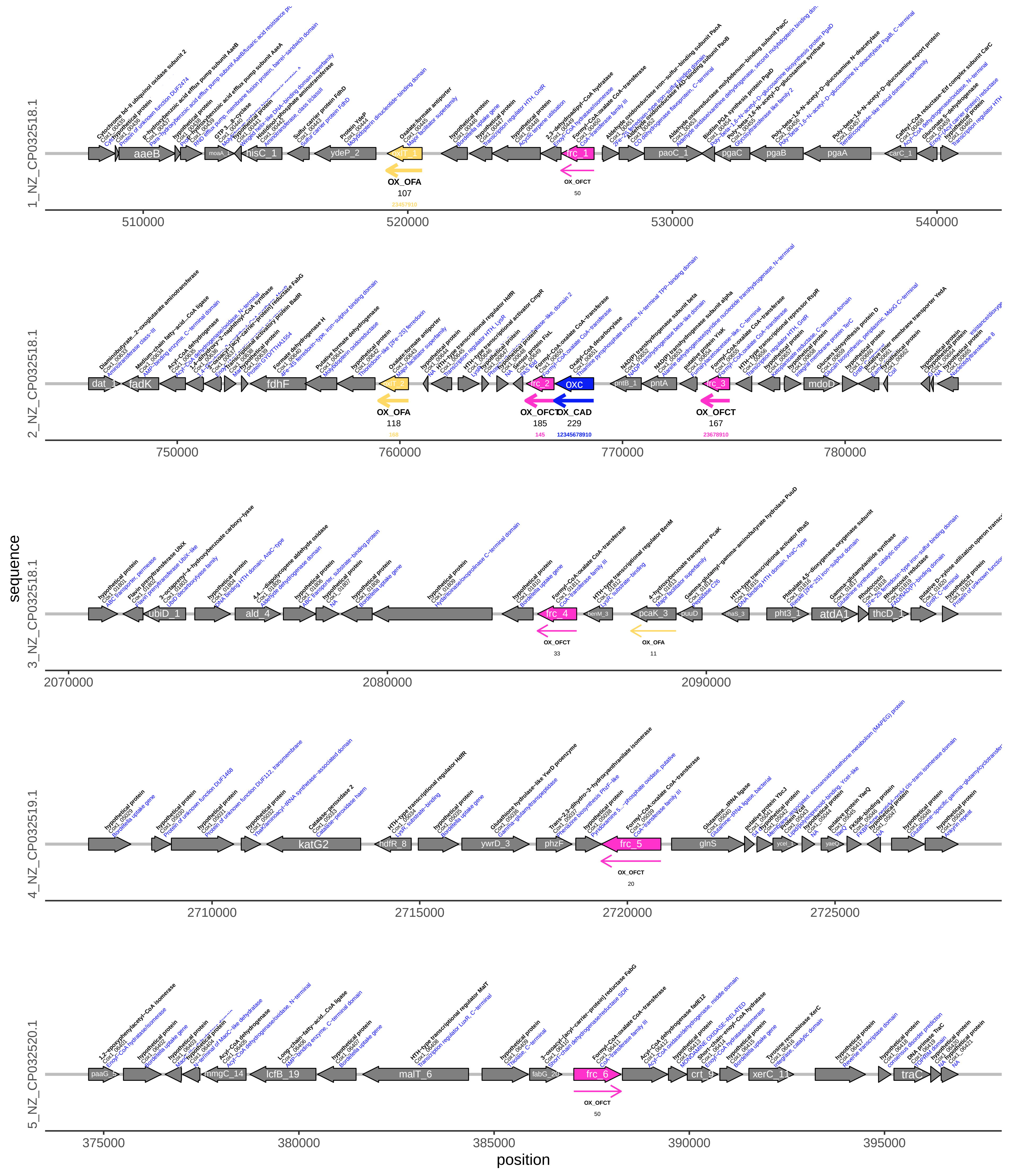

### Cox2.pdf

#### Cox2 (GCF_004768545): Cupriavidus oxalaticus X32 (fabio)

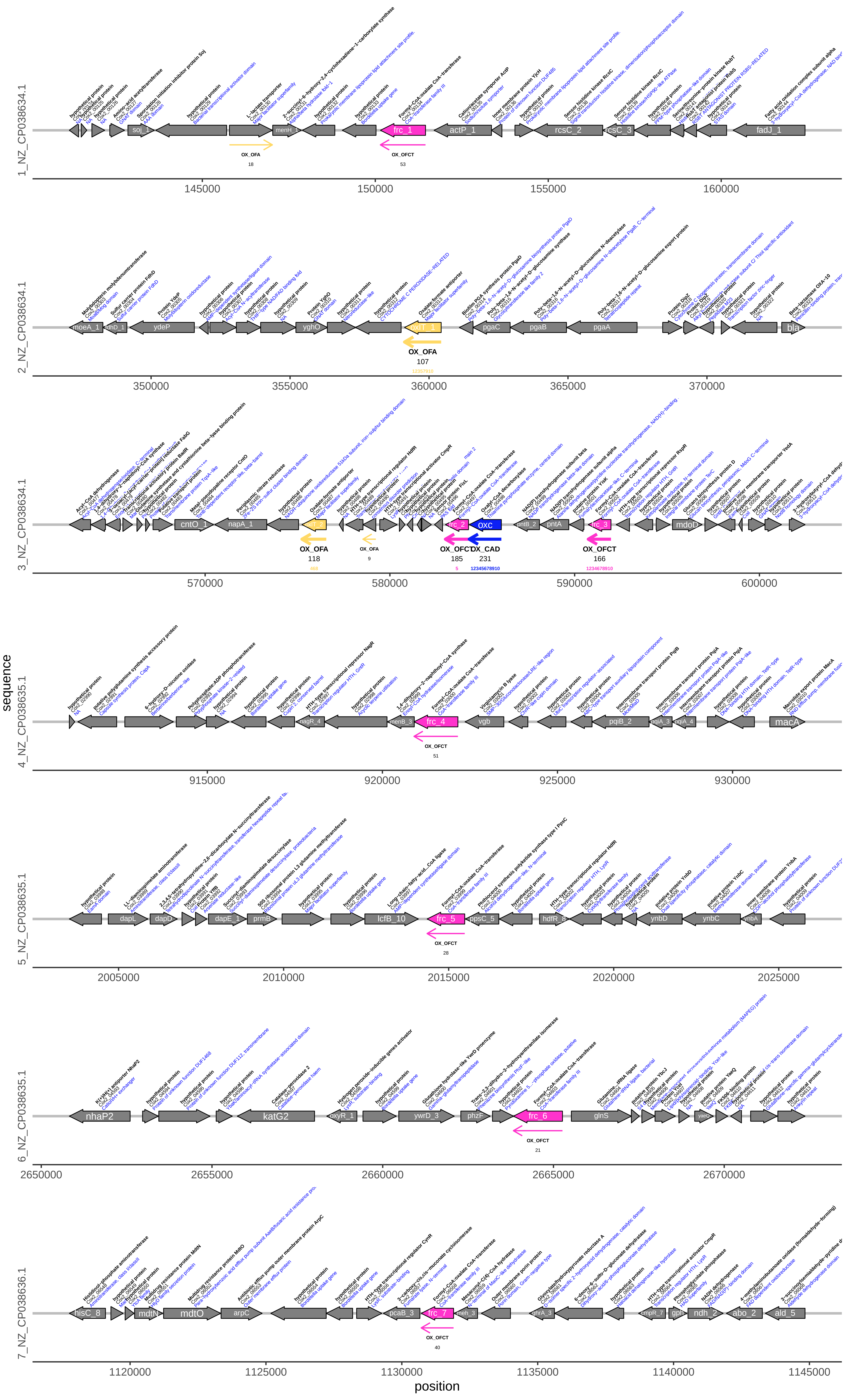

### Mmas.pdf

#### Mmas (MGYG000002310): Microvirga massiliensis (gut)

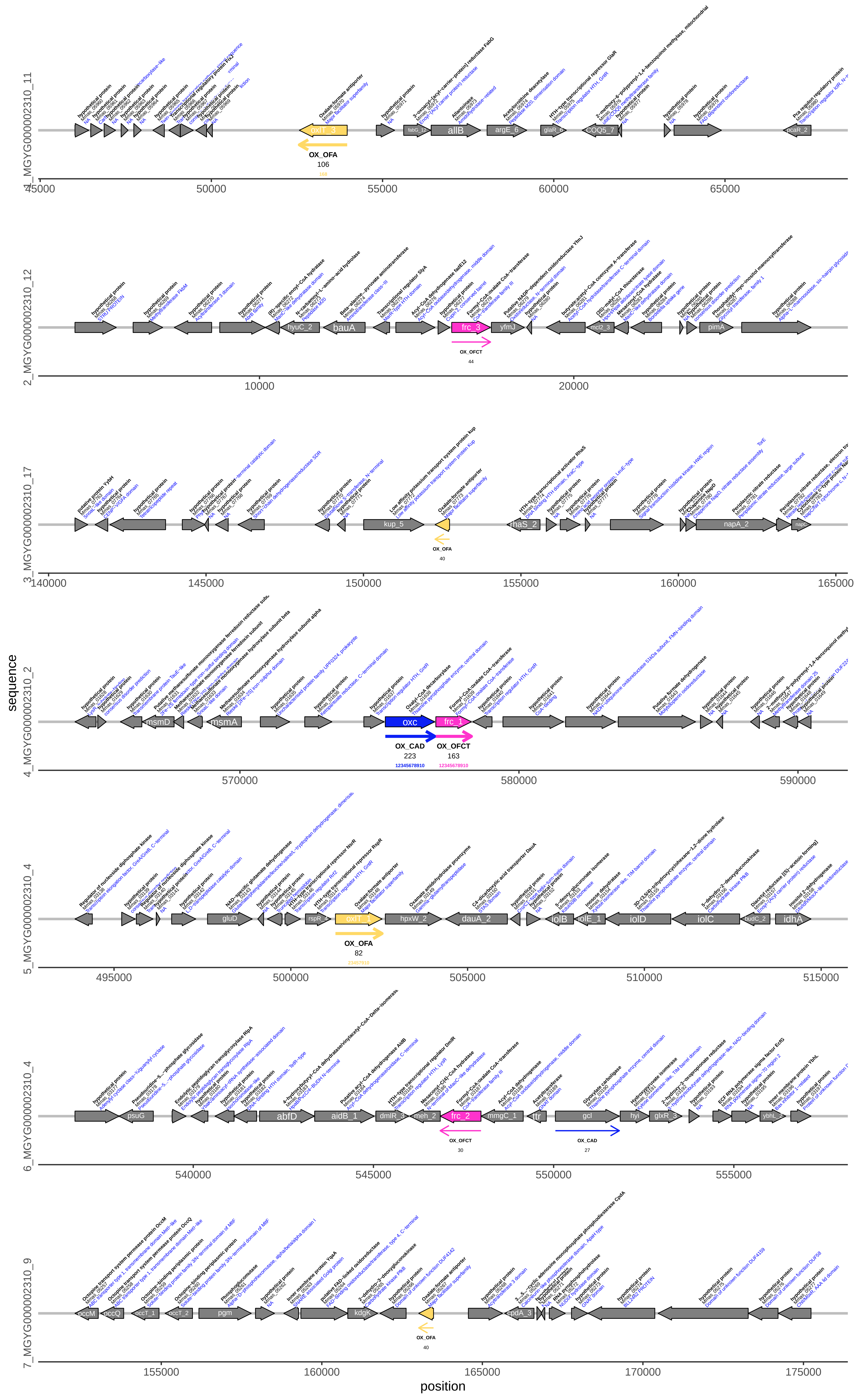

### Msp1.pdf

Msp1 (MGYG000001265): Methylobacterium sp002778835 (gut)

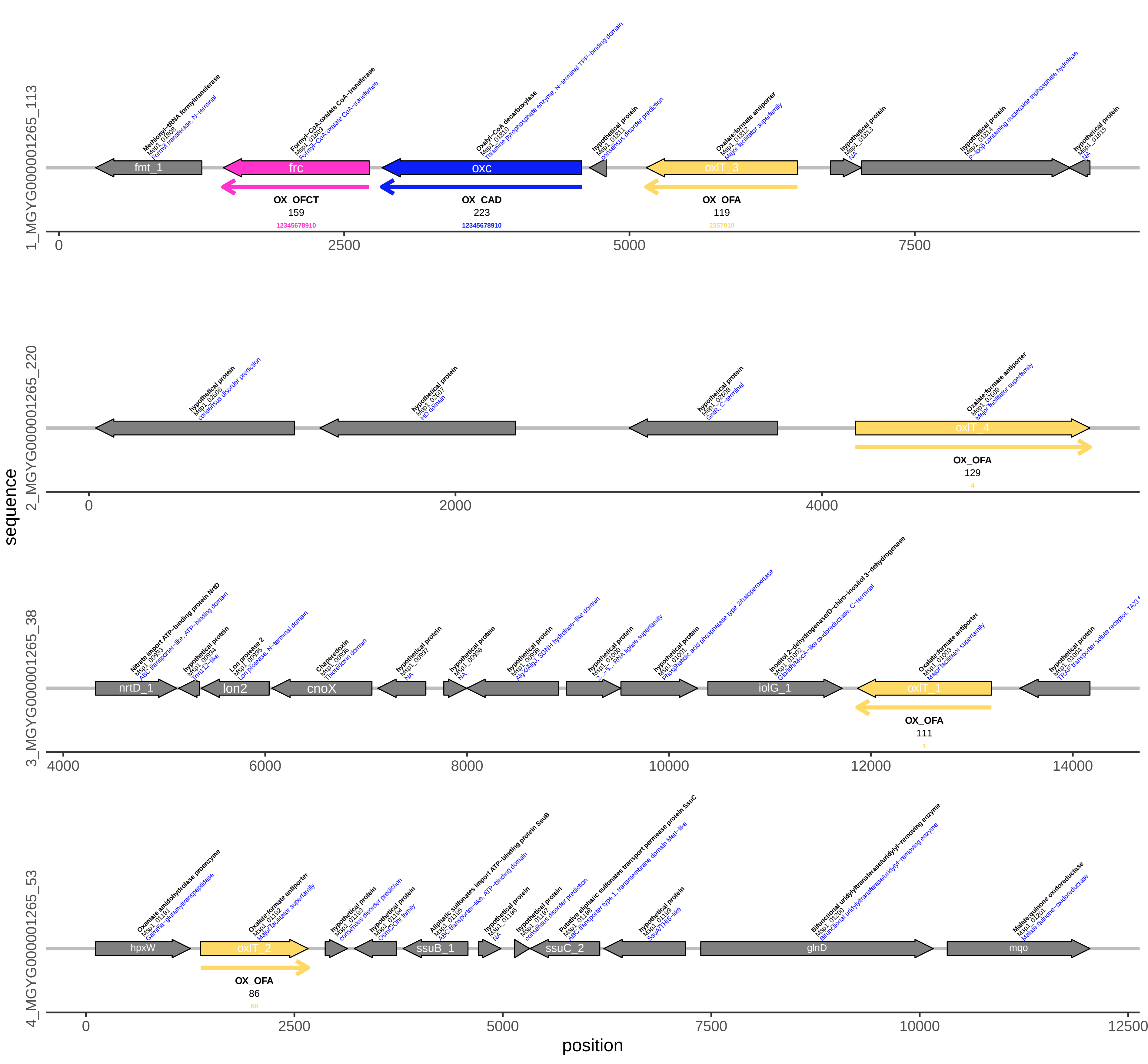

### Ofo1.pdf

#### Ofo1 (MGYG000001331): Oxalobacter formigenes (gut)

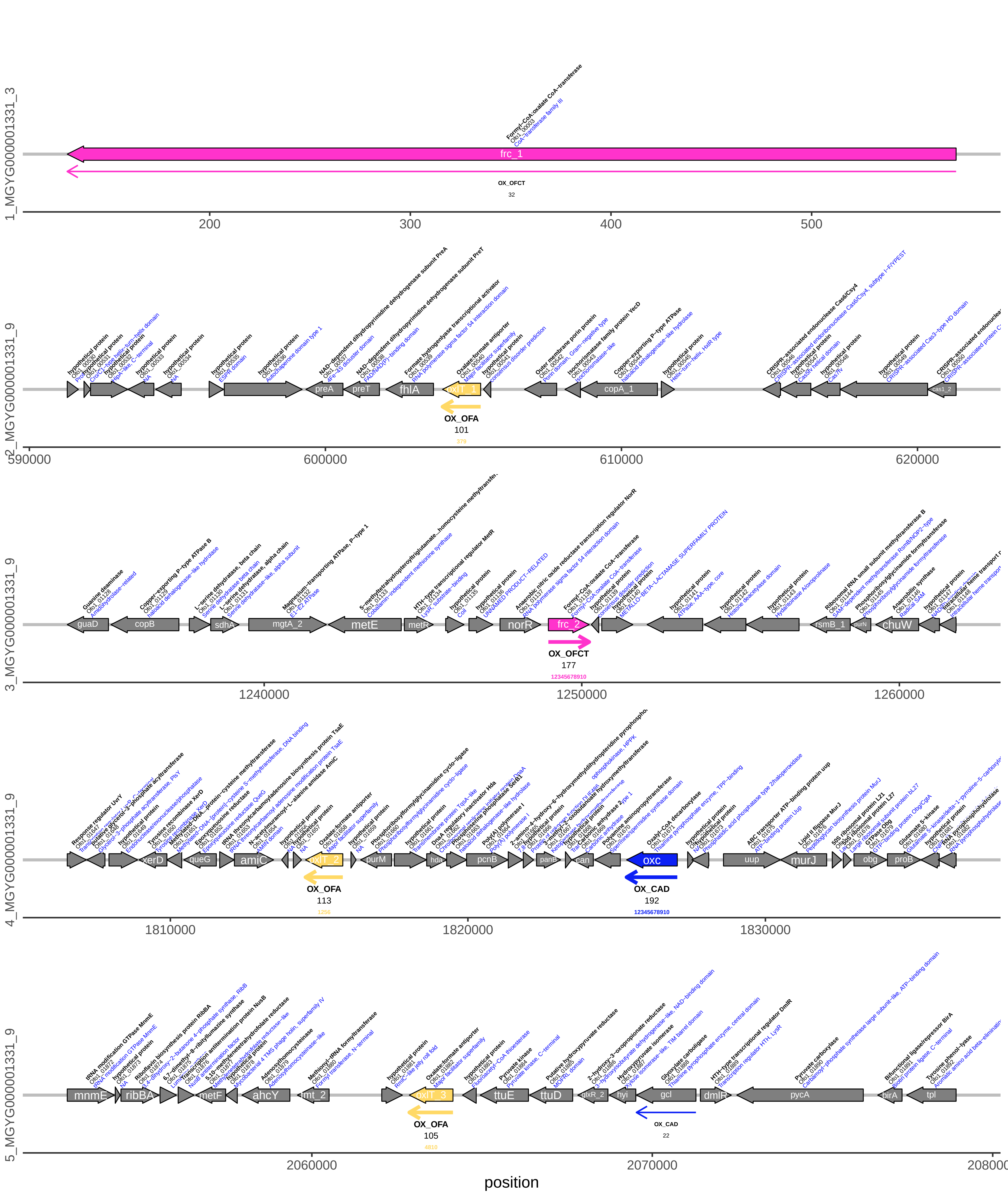

### Ofo2.pdf

#### Ofo2 (GCF_002128345): Oxalobacter formigenes OXCC13 (fabio)

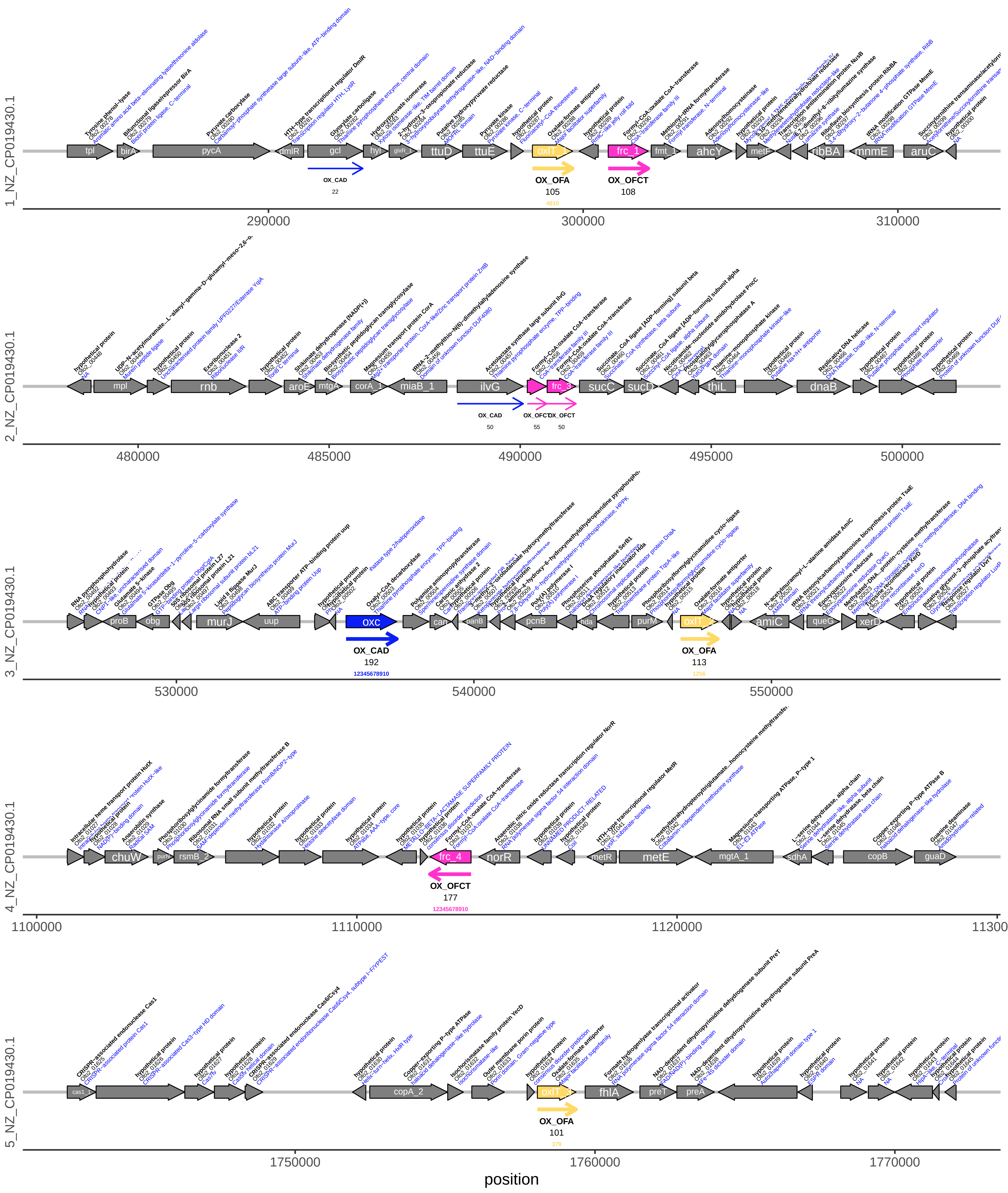

### Ofo3.pdf

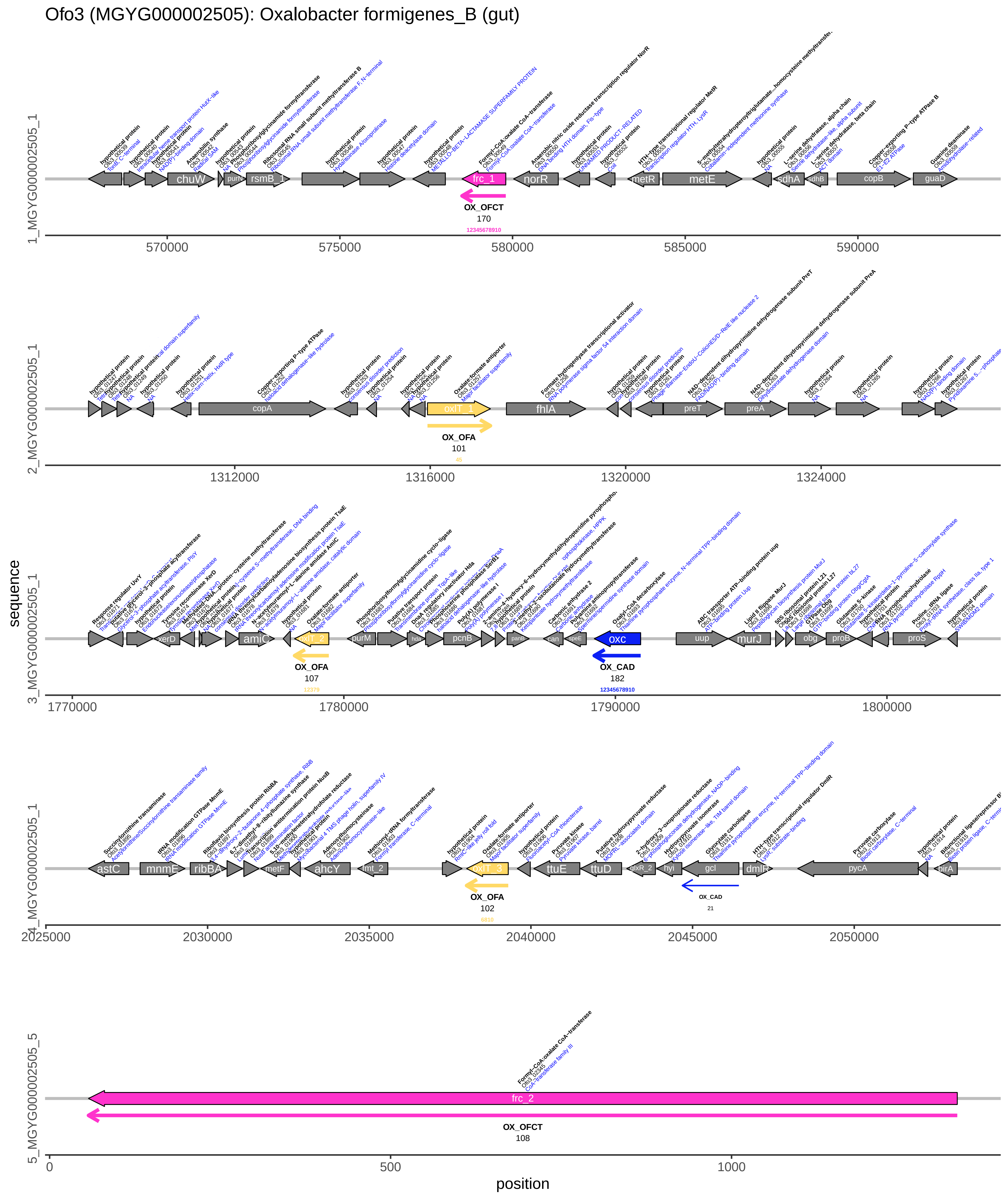

### Osp1.pdf

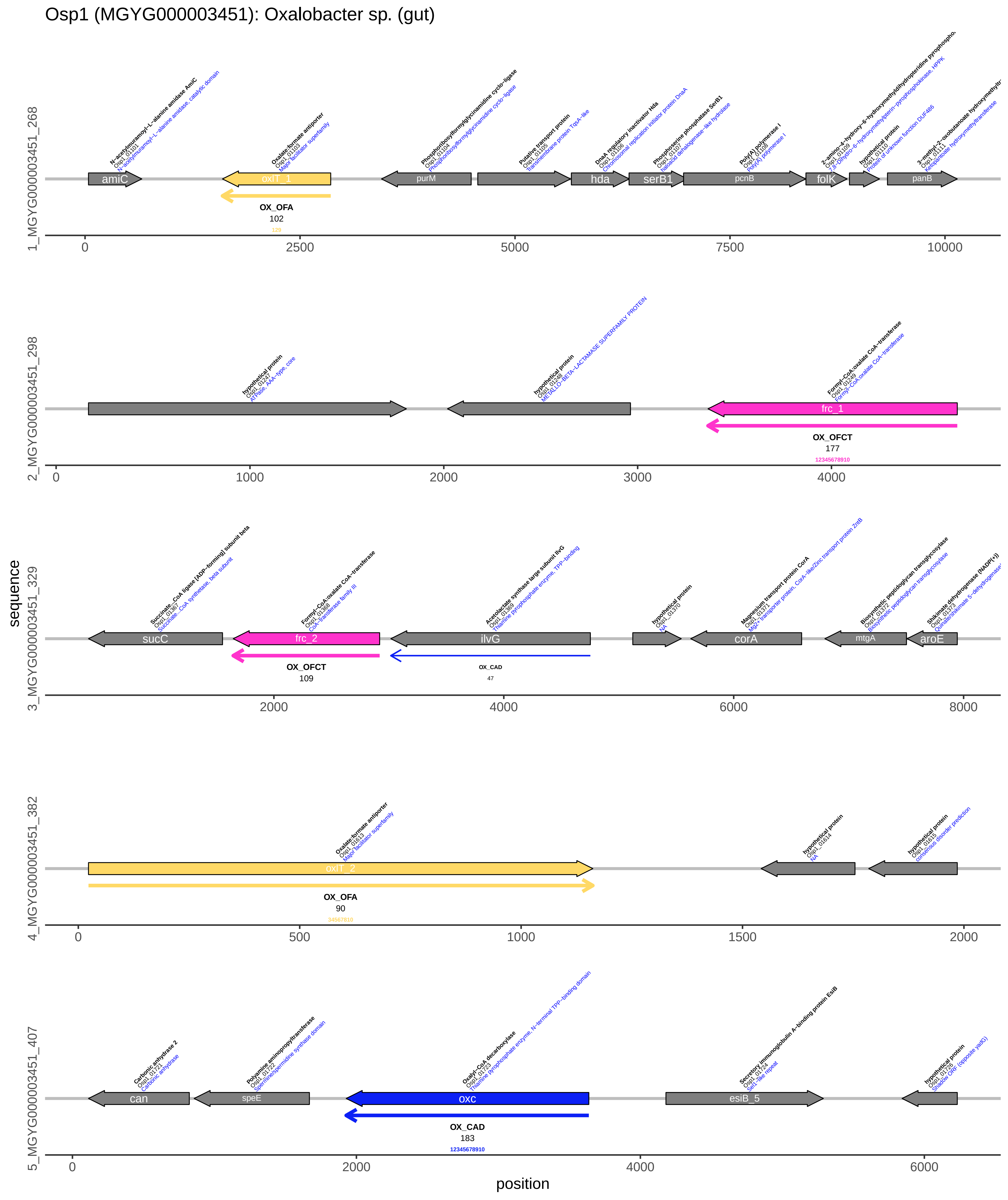

### Osp2.pdf

#### Osp2 (MGYG000004063): Oxalobacter sp900760095 (gut)

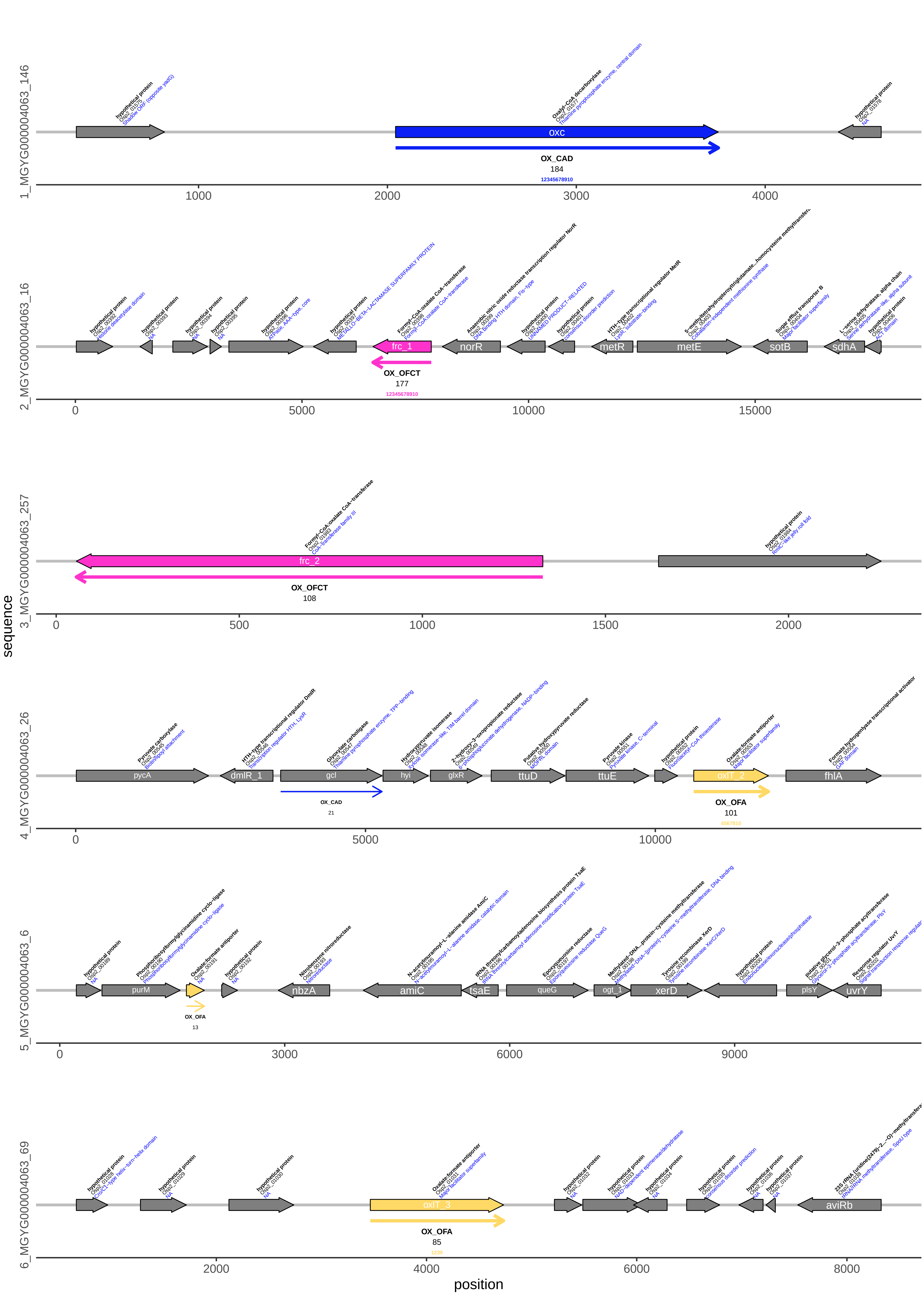

### Osp3.pdf

Osp3 (MGYG000002703): Oxalobacter sp905202055 (gut)

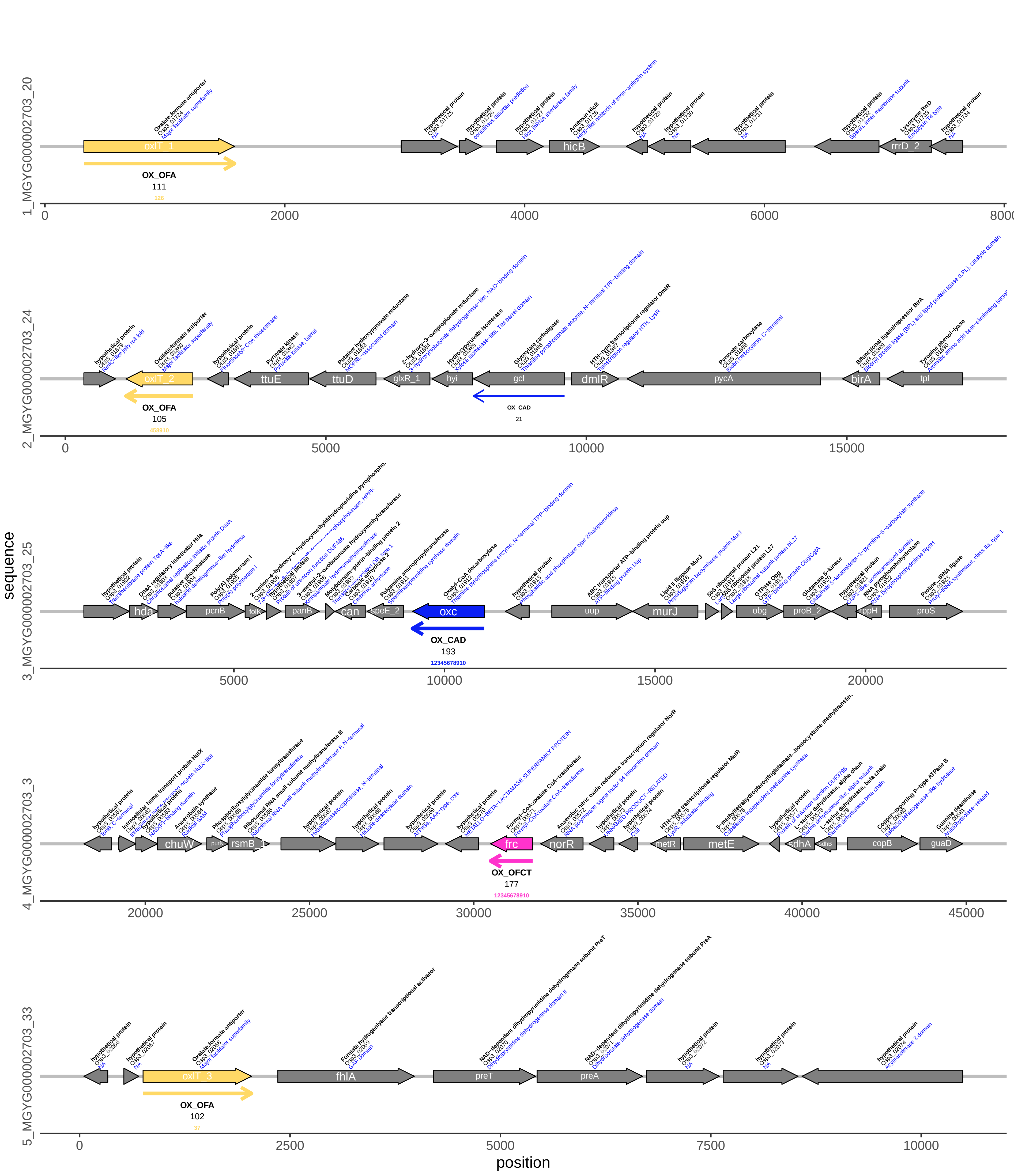

### OX_CAD-maxrn-ro-scicol-nolbl.png

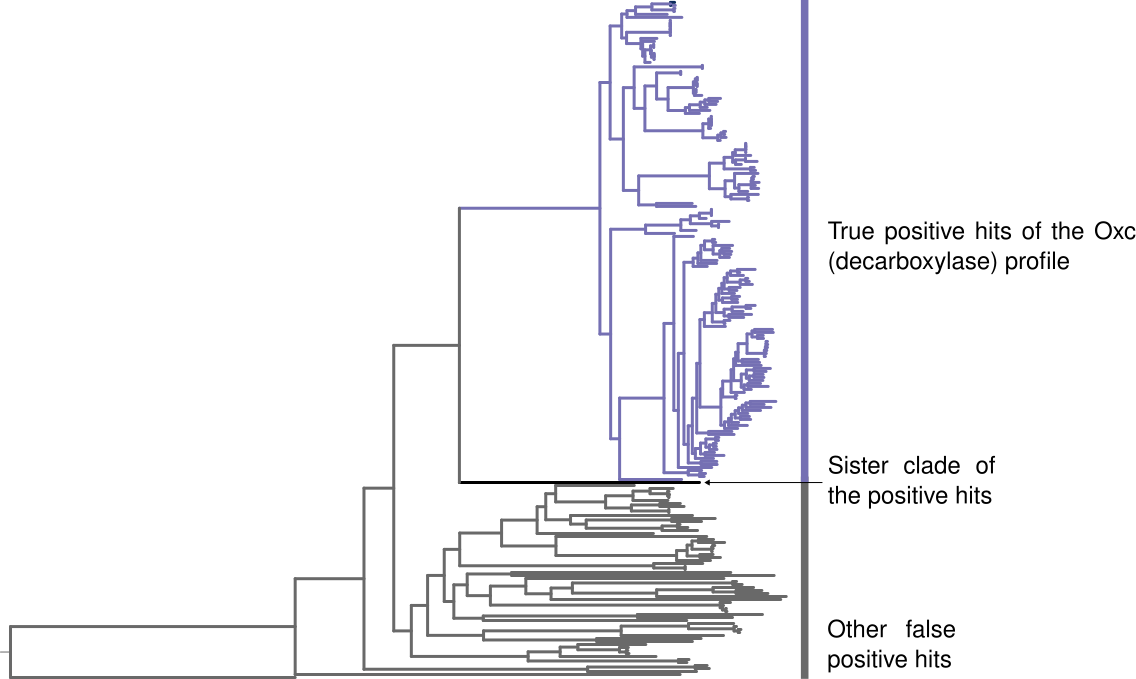

### OX_CAD_highlit_TP_hmap-binary-avg.pdf

OX_CAD, sampled matches, true positives clade

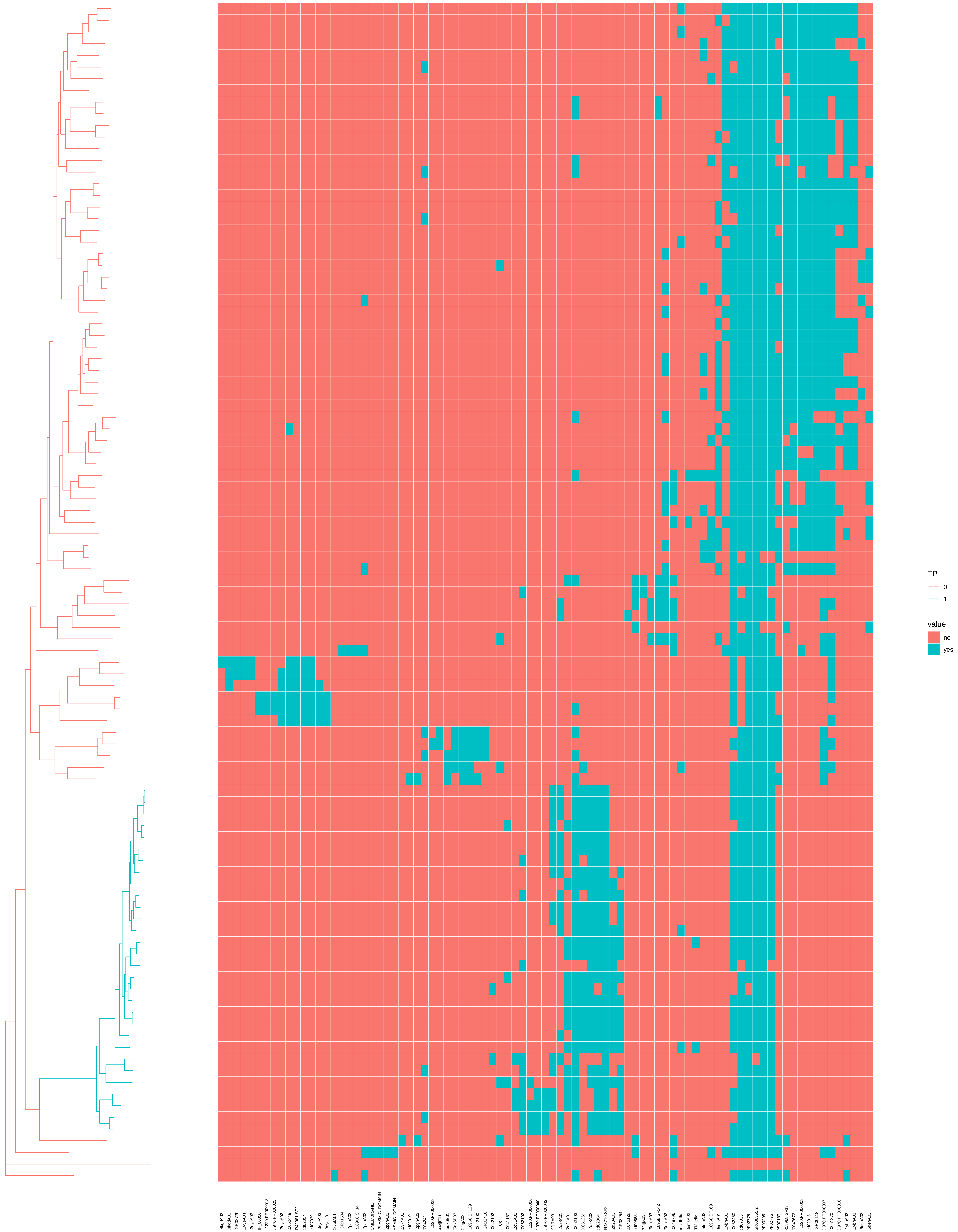

### OX_CAD_highlit_TP_hmap-euclidean.pdf

OX_CAD, sampled matches, true positives clade

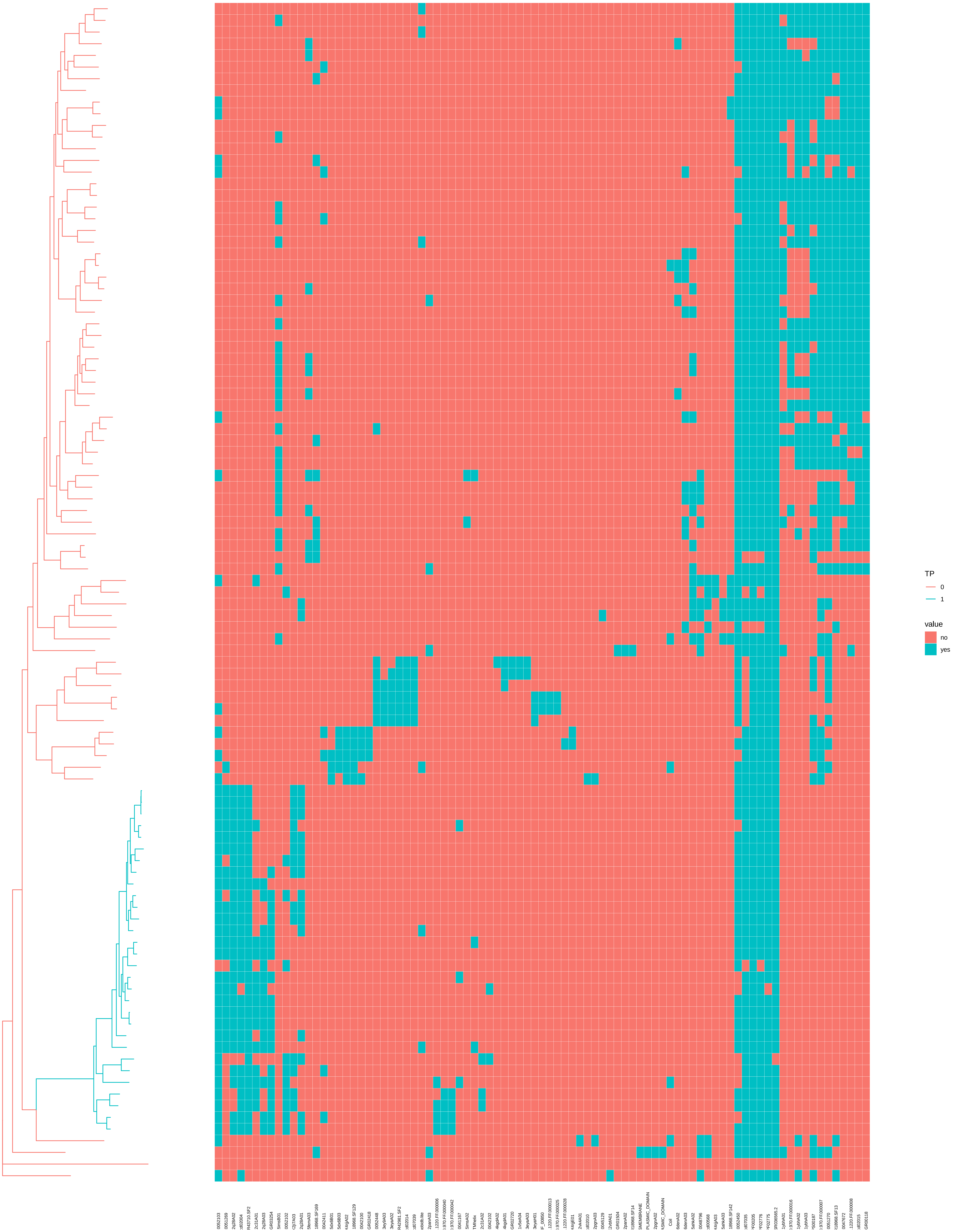

### OX_OFA-maxrn-ro-scicol-nolbl.png

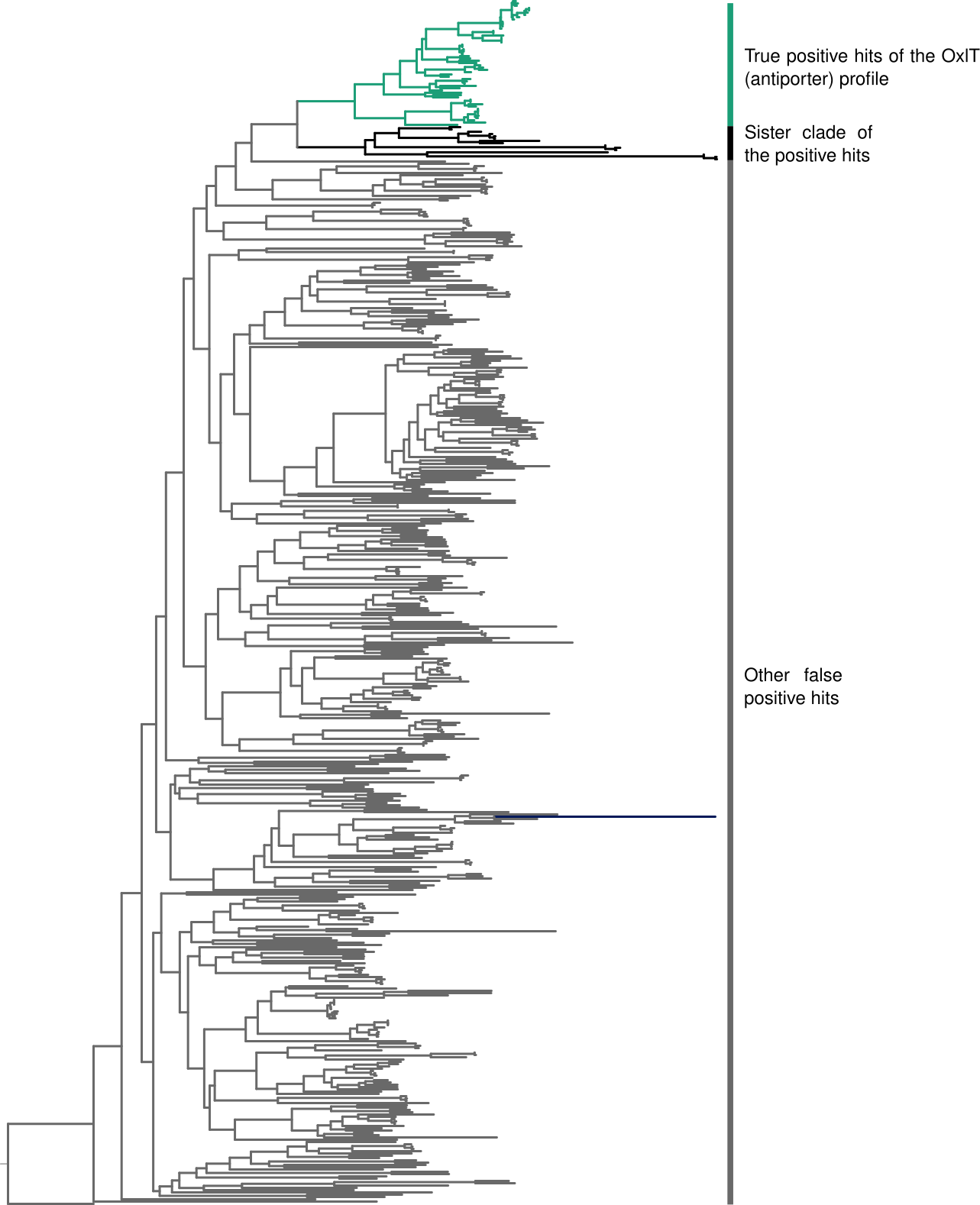

### OX_OFA-pg2.msa.png

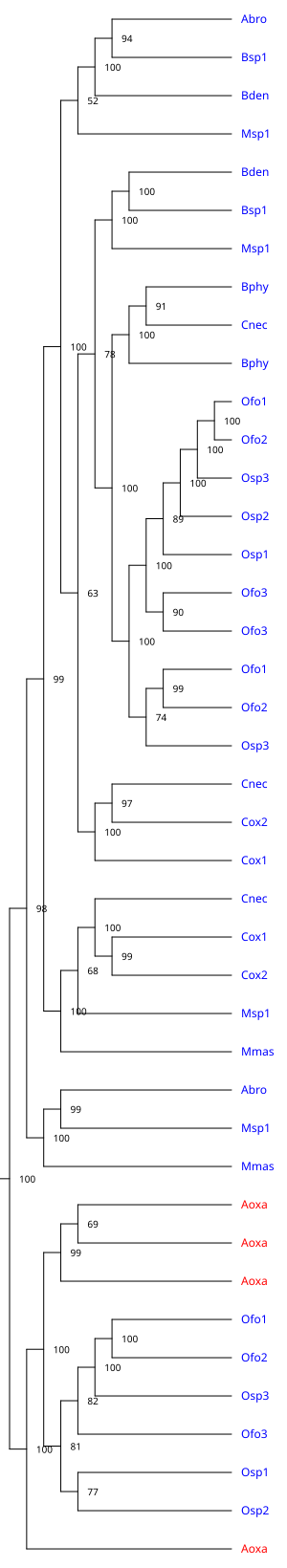

### OX_OFA_highlit_TP_hmap.pdf

OX_OFA, sampled matches, true positives clade

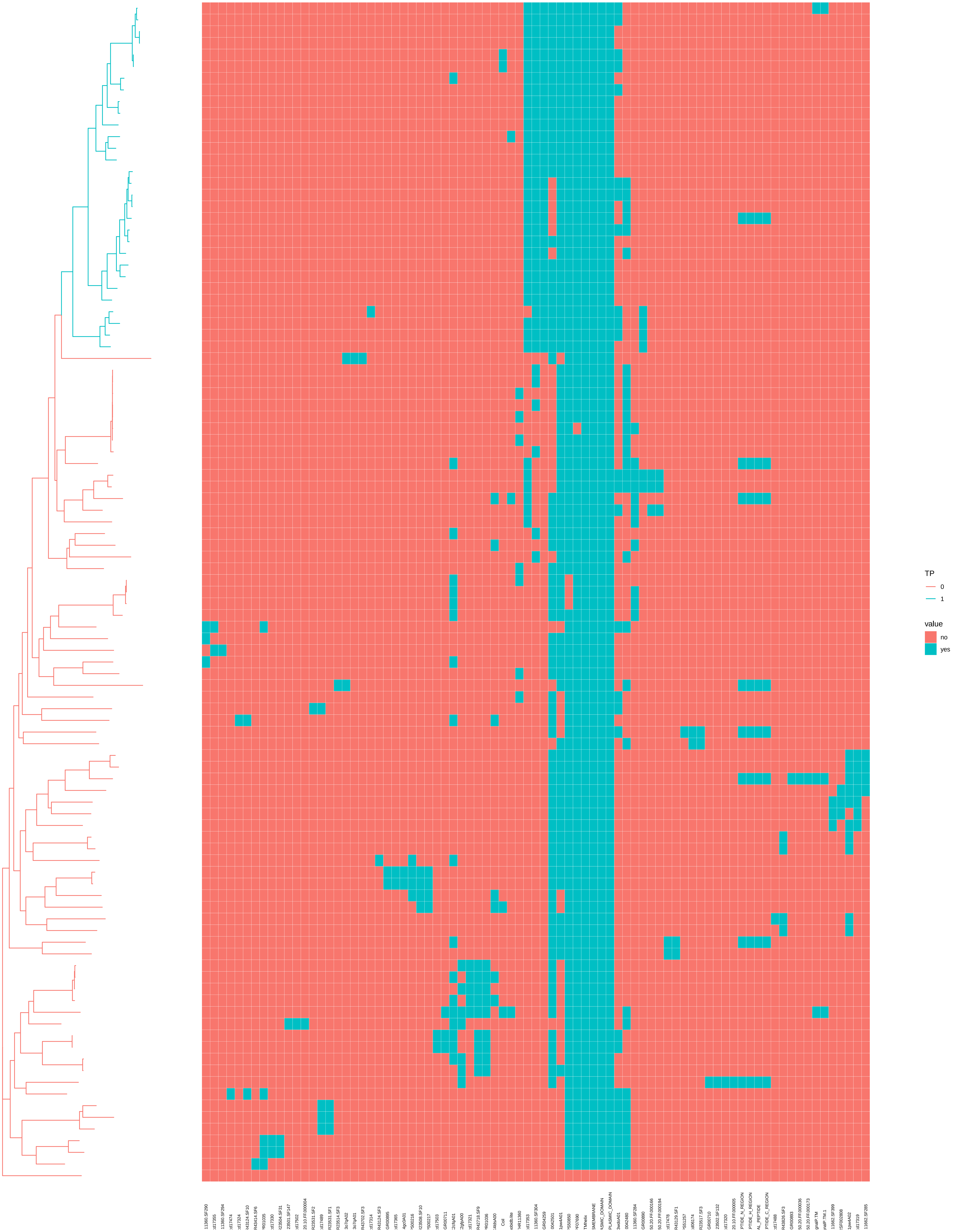

### OX_OFCT-maxrn-ro-scicol-nolbl.png

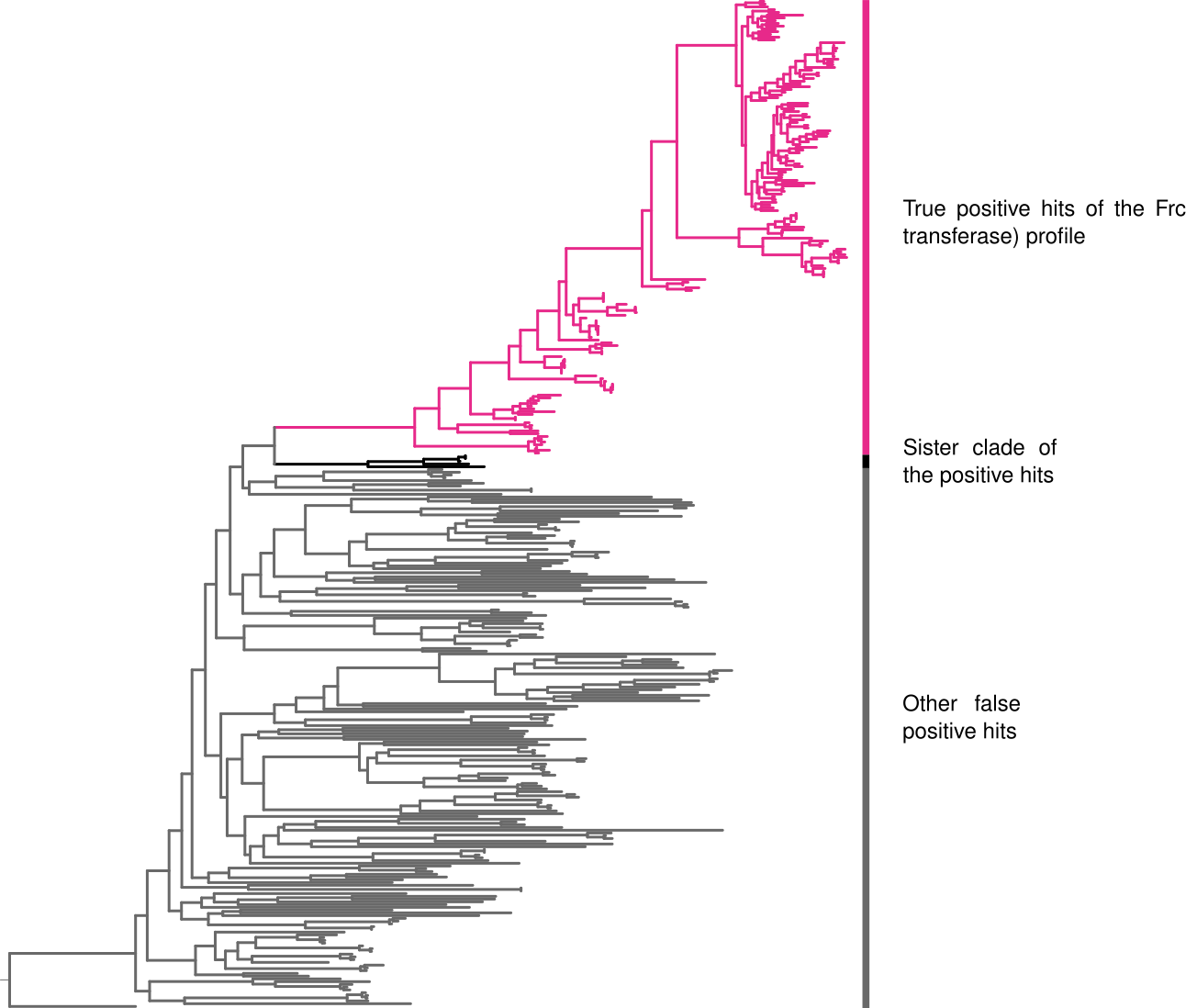

### OX_OFCT_highlit_TP_hmap.pdf

OX_OFCT, sampled matches, true positives clade

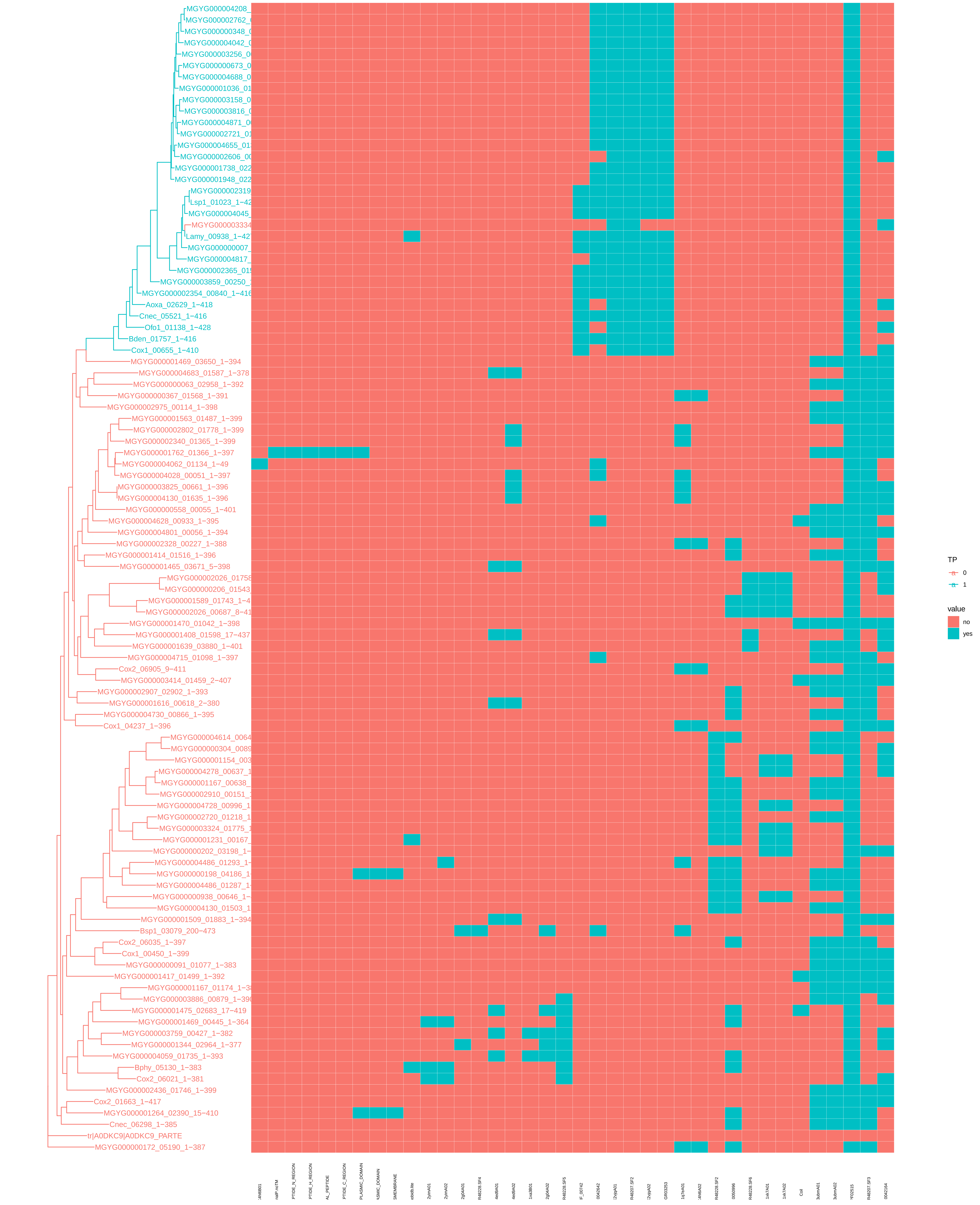
